# Nanoscale remodeling of mitochondrial architecture during mtRNA surveillance failure

**DOI:** 10.64898/2026.10.20.759735

**Authors:** Eve V. Kakudji, Kevin Rose, Jaewon Lee, Eric Herrmann, Priya Gatti, James H. Hurley, Samantha C. Lewis

## Abstract

Mitochondria within a cell exhibit heterogeneity in their composition and metabolic functions, and how this specialization shapes the responses of individual organelles to stress is unclear. We applied fluorescence microscopy and *in situ* cryo-electron tomography to examine the composition of mitochondrial stress bodies and their relationship to inner mitochondrial membrane architecture in the context of a mitochondrial genome expression defect. We discovered that defective mitochondrial RNA processing prompted a loss of inner membrane domain identity, leading to the mislocalization of ATP synthase from cristae edges to the boundary membrane, perturbed cristae morphometrics, and complete cristae loss in a subset of organelles. Mitochondrial stress bodies were enriched with nucleic acid and protein components linked to oxidative pathways, while separate and distinct mitochondrial domains became enriched with oligomerized P5CS, a marker of reductive proline synthesis. We found that consolidation of mitochondrial RNA into stress bodies, as well as ATP synthase remodeling, were regulated independently from P5CS concentration into a subset of the mitochondria. We propose that stress compartmentalization enables independent tuning of the quality control of oxidative pathway components via spatial sequestration from the rest of the mitochondrial network.

## INTRODUCTION

Mitochondria perform diverse functions in eukaryotic cells, including energy production, amino acid synthesis, and the maintenance of cellular ion and phospholipid homeostasis. They also harbor their own polyploid genome, whose expression is integrated with mitochondrial bioenergetic functions (1,2). Within each cell, mitochondria form a dynamic network that responds to metabolic signals and local energy demands. Because mitochondrial dysfunction contributes to metabolic and neurodegenerative diseases, understanding how mitochondrial genome regulation is integrated with these subcellular cues is likely to provide insight into etiology and pathology of human disease. However, how mitochondrial DNA (mtDNA) synthesis, distribution, and expression vary within individual mitochondria, and how these processes are spatially remodeled during disease or stress states, remains poorly understood.

The processing and eventual degradation of polycistronic mitochondrial RNA (mtRNA) encoded in the mitochondrial genome are critical for maintaining mitochondrial function and regulation of innate immunity (3,4). Homeostatic mitochondrial RNA degradation elides the accumulation of immunogenic double-stranded mitochondrial RNA (mt-dsRNA), that forms via complementary binding of mtRNA species transcribed from opposite mtDNA strands. The ATP-dependent RNA helicase SUV3 (SUPV3L1) and the mitochondrial polynucleotide phosphorylase (PNPase) complex function in a linear pathway to unwind and degrade mtRNA in cells (5). Disruption of this pathway allows mt-dsRNA to accumulate and, when released into the cytosol, activate antiviral type I interferon and proinflammatory signaling (6–8).

We recently described mitochondrial stress bodies (MSBs), a previously unrecognized layer of post-transcriptional regulation of mitochondrial biogenesis and stress responses in human cells (9). We found that a subset of mitochondria within cells preferentially support electron transport chain (ETC) biogenesis through localized translation of mtDNA-encoded mRNAs, which we termed translation hubs (9). These mitochondrial translation hubs are particularly sensitive to RNA-processing stress, which prompts their conversion to matrix protein aggregates that sequester mitochondrial messenger RNA, functional RNA, and mt-dsRNA (9,10). Concurrently, mitochondrial translation was selectively suppressed, as has been observed in other cell stress contexts including hypoxia and ethidium bromide exposure (9–12).

Beyond mtDNA expression, mitochondrial biosynthetic pathways are likewise spatially organized in cells (13–17). Pyrroline-5-carboxylate synthase (P5CS), a key enzyme in proline and ornithine biosynthesis, converts glutamate to glutamate-5-semialdehyde (GSA), which spontaneously cyclizes to pyrroline-5-carboxylate (P5C). P5CS oligomerizes into filaments during high bioenergetic demand, but redistributes upon supplementation with proline or ornithine (18). Oligomerization of P5CS into a subset of mitochondria requires cycles of mitochondrial fusion and fission, and presumably content mixing and cristae reorganization.

Cristae morphology serves as a physical barrier to the movement of molecules and is linked to the ability of mitochondria to be specialized. Cristae can adopt a variety of morphologies depending on cell type, and are constitutively remodeled during mitochondrial fusion and fission cycles that include intra-mitochondrial partitioning, merging, or vesicularization of the inner membrane (19–22). Perturbed cristae morphometrics are a common feature of human cancer and neurodegenerative disease (23–27).

Mitochondrial ultrastructure also shapes stress responses more directly by regulating the organization and/or integrity of the mitochondrial membranes that prevent mt-dsRNA leakage to the cytosol in homeostasis, protein folding or misfolding that can make specific subsets of organelles preferential targets for mitophagic turnover, as well as the gene expression machinery needed for stress recovery (28–32). Disrupted mitochondrial proteostasis activates protective pathways, including the mitochondrial unfolded protein response (UPRmt), which responds to matrix protein misfolding through local mitochondrial changes and nuclear transcriptional programs (33–35). These responses suppress protein translation and increase the expression of matrix proteostasis factors, including the canonical HSP60 chaperone complex and AFG3L2 (29,30,36–39). These observations suggest that mitochondrial metabolic enzymes, RNA-processing and translation machinery, all undergo coordinated spatial reorganization in crisis.

Here, we combine fluorescence microscopy and *in situ* cryo-electron tomography to investigate how stress-induced mitochondrial remodeling spatially compartmentalizes the oxidative and reductive biosynthetic machineries into distinct network compartments. We find that the oxidative machinery associated with mitochondrial translation and RNA processing is sequestered into MSB-positive mitochondria, whereas the reductive enzyme P5CS occupies a distinct, non-overlapping mitochondrial subset within the same cells. This compartmentalization is accompanied by matrix protein aggregation and inner membrane remodeling, including redistribution of ATP synthase complexes from cristae to the inner boundary membrane and subsequent cristae vesicularization/collapse that cloisters gene expression machinery from P5CS filaments even within the shared outer membrane of individual mitochondria. We propose that this spatial compartmentalization serves to enable differential regulation of oxidative and reductive biosynthetic pathways during bioenergetic stress by spatially separating these components.

## RESULTS

### Mitoribosome components are sequestered in mitochondrial stress bodies

The machineries of mitochondrial RNA transcription, processing and translation are particularly sensitive to proteotoxic stress caused by inhibition of HSP90 complexes, as compared to the general matrix proteome (33). Thus, we sought to determine whether mitoribosomal components may be sequestered within MSBs, in addition to mt-RNA species and mitochondrial RNA binding proteins (RBP). We induced the formation of translationally-inhibited mitochondrial stress bodies (MSB) in U2OS human osteosarcoma cells via depletion of the SUV3 helicase (**Fig. 1A**), then examined the micron-scale distribution of MRPL23, a mitoribosomal component whose solubility is highly responsive to pharmacological inhibition of HSP90 (33).

**Figure 1:**
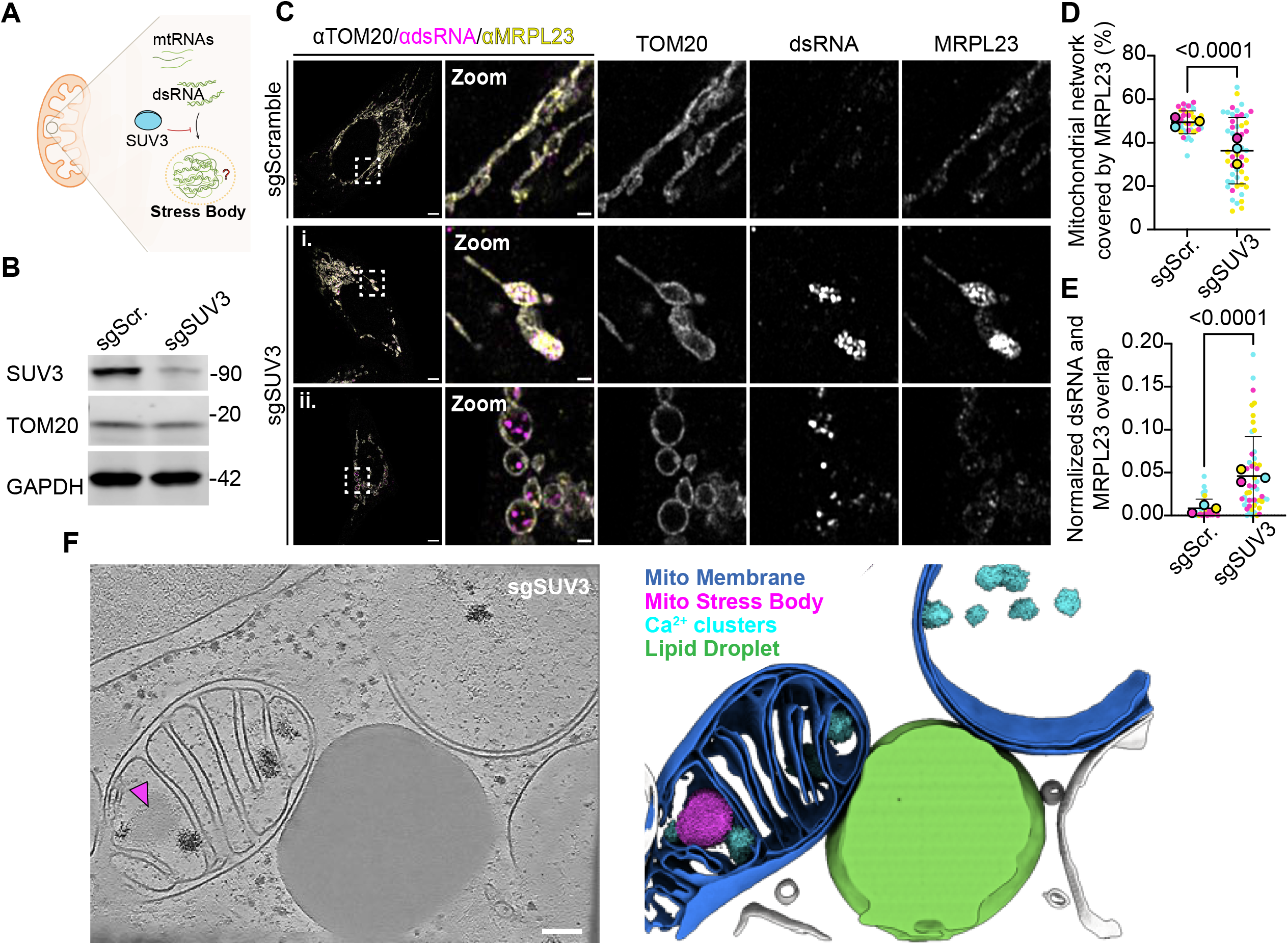
MRPL23 is reorganized and recruited to dsRNA bodies under sgSUV3 stress.(A) Schematic illustrating mitochondrial stress body formation following sgSUV3 stress **(B)** SUV3, TOM20, and GAPDH protein levels in sgScramble versus sgSUV3 U2OS-Cas9 cells analyzed by western blot. GAPDH is used as a loading control. **(C)** Immunofluorescence images of U2OS-Cas9 transfected with sgScramble (top) and sgSUV3 (bottom), and immunolabeled with anti-TOM20, anti-dsRNA (J2), and anti-MRPL23. Imaged 7 days after transfection. Scale bars: 5 μm; 1 μm in zoom. A single slice of a z-stack is shown for each condition. **(D)** Quantification of the experiment performed in (C) showing MRPL23 area covering TOM20-segmented mitochondrial network. Quantification performed on maximum intensity projection images. **(E)** Quantification of the experiment performed in (C) showing the Jaccard Index, labeled as overlap, and defined as the sum area of MRPL23 and dsRNA overlapping over the sum area of unique MRPL23 and dsRNA. Quantification performed on maximum intensity projection images. For (D), N = 44 cells (sgScramble) and 52 cells (sgSUV3), and for (E), N = 45 cells (sgScramble) and 50 cells (sgSUV3) over three independent biological replicates. Error bars display mean values ± standard deviation, and the bottom bar is clipped if below 0. Statistical analysis performed on individual data points with Mann-Whitney test. (**F**) Cryo-ET tomogram of sgSUV3-transfected U2OS cells and corresponding 3D segmentation depicting an example of a mitochondrion with an aggregate. Scale bar: 100nm.

We depleted SUV3 from U2OS cells by CRISPR-Cas9 technology using three guide RNAs targeting exon 1, and visualized the distribution of MRPL23, TOM20, and dsRNA using indirect immunofluorescence (**Fig. 1B-C**). In control cells, MRPL23 was evenly distributed throughout the extensive mitochondrial network (**Fig. 1C**). In contrast, MRPL23 coalesced with dsRNA into MSBs in SUV3-depleted cells, consistent with the sequestration of the mitoribosomal components within this subset of mitochondria (**Fig. 1Ci**). Three-dimensional segmentation of MRPL23 and dsRNA within the stress bodies suggests a structured organization in which the mitoribosomal protein is scaffolded around dsRNA (**Fig. S1**). Consistently, spatial analysis revealed a significant reduction in the proportion of total mitochondria that contained detectable MRPL23 (**Fig 1D**), as well as a concurrent increase in MRPL23-dsRNA colocalization (**Fig. 1E**). However, this colocalization was dependent on dsRNA density (**Fig. 1Cii**).

To better visualize the nanoscale features of individual mitochondria during MSB formation and MRPL23 sequestration, we optimized SUV3-depleted U2OS cells for *in situ* cryo-electron tomographic analysis (cryo-ET). We performed cryo-FIB milling of sgSUV3 cells to generate lamellae of approximately 200 nm thickness for ultrastructural study (**Fig. S2**). We reconstructed 34 tomograms, and segmented mitochondrial membranes as well as other organelles in contact with mitochondria such as lipid droplets, and the MSB within the mitochondrial matrix (**Fig. 1F**).

MSB occupancy varied between individual mitochondria, consistent with the fluorescence microscopy, as did cristae density. We noted that regularly-spaced, prominent cristae coincided with the presence of at least one MSB, while in this case mitochondria lacking MSB also lacked cristae altogether (**Fig. 1F**). These data highlight a potential role for MSB as a protective reservoir for mitochondrial proteins and RNA during proteotoxic stress, and led us to quantitatively profile cristae architecture in SUV3-depleted cells, as described below.

### Mitochondrial stress body formation is associated with cristae remodeling

Previous work has implicated aggregates of misfolded mitochondrial proteins in cristae deformation, eventually leading to their loss (29). However, our observations of intact transverse cristae within the subset of mitochondria that contain MSB led us to consider that, in the context of mitochondrial RNA processing stress, aggregates could serve a protective role by, for example, sequestering disassembled mitoribosomes, partially processed RNA, and/or misfolded proteins from the broader mitochondrial network.

To quantitatively examine the ultrastructural consequences of SUV3 depletion on cristae, we performed cryo-ET to visualize mitochondrial ultrastructure at nanoscale-resolution. While control cells primarily exhibited straight cristae, SUV3-depleted cells contained mitochondria with a variety of cristae morphologies, including straight, collapsed, and vesicular cristae (**Fig. 2A-B**). Strikingly, SUV3-depleted cells also contained another subset of mitochondria that were devoid of cristae altogether, in which the inner membrane was completely parallel to the outer membrane (**Fig. 1F**, **Fig. 2A-B**). To validate that these double-membrane structures were indeed mitochondria, we complimented this analysis with live 3D Airyscan microscopy of SUV3-depleted cells using the mitochondrial inner-membrane stain Mitotracker Green. Live-cell imaging indeed revealed a subset of swollen mitochondria with smooth interior Mitotracker Green signal, suggesting a simple IMM organization relative to control cells (**Fig. S3**).

**Figure 2:**
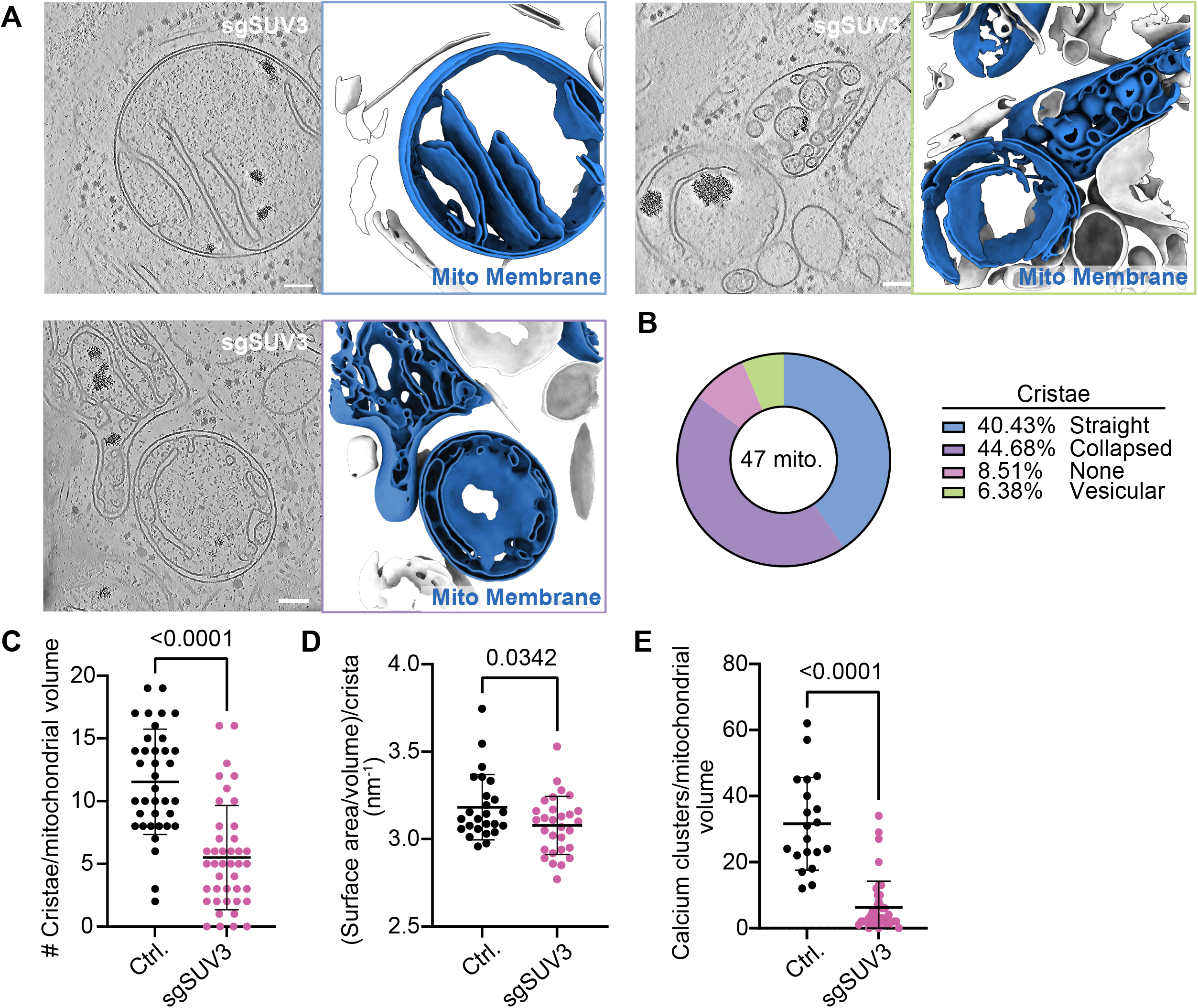
**sgSUV3 stress induces cristae remodeling.(A**) Cryo-electron tomography images of sgSUV3-transfected U2OS cells and corresponding 3D segmentations. Scale bar: 100nm. (**B)** Proportion of mitochondrial cristae phenotypes observed. N = 47 mitochondria. **(C)** Quantification of cristae density per mitochondrial volume in control U2OS and U2OS cells transfected with sgSUV3. N= 37 control mitochondria, N = 42 sgSUV3 mitochondria. **(D)** Cristae surface area and volume compared between control and sgSUV3-transfected cells. N=25 independent control cristae, N=29 independent sgSUV3 cristae. **(E)** Quantification of calcium clusters per mitochondrial volume in control and sgSUV3-transfected cells. N =20 control mitochondria, N=42 sgSUV3 mitochondria. Error bars display mean values ± standard deviation, and the bottom bar is clipped if below 0. Statistical analysis performed using an unpaired t-test applied to the represented data points.

We found that the overall density of detectable cristae segmented per mitochondrial volume was significantly decreased in SUV3-depleted cells (**Fig. 2C**). We also found that cristae contraction, as reported by the ratio of membrane surface area to volume per isolated crista, was reduced within SUV3-depleted cells (**Fig. 2D**), as were electron-dense mitochondrial calcium clusters (**Fig. 2E**). Overall, these data demonstrate that MSB formation downstream from defective mtRNA quality control is a previously unappreciated condition associated with mitochondrial cristae remodeling.

### Mitochondrial stress body formation is a common response to mitochondrial dysfunction

We next wondered whether MSB formation and the associated cristae phenotypes were specific to SUV3 depletion or instead generalizable to other modes of mitochondrial dysfunction. Specifically, we considered the potential reorganization of mitochondrial RNA under inhibition of electron transport chain (ETC) components using oligomycin and antimycin A (OA). To test this, we transfected U2OS cells with a plasmid for transient transfection encoding an mCherry-tagged isoform of human TOM20, applied the live-cell mitochondrial RNA stain RNASelect, and performed live-imaging of OA-treated cells. Live imaging revealed the concentration of mitochondrial RNA within the swollen mitochondria of some of the OA-treated cells (**Fig. 3A**). In contrast, RNA was sparingly distributed along the tubular network of untreated cells (**Fig. 3A**). These results indicate that mitochondrial depolarization is associated with internal mitochondrial RNA reorganization.

**Figure 3:**
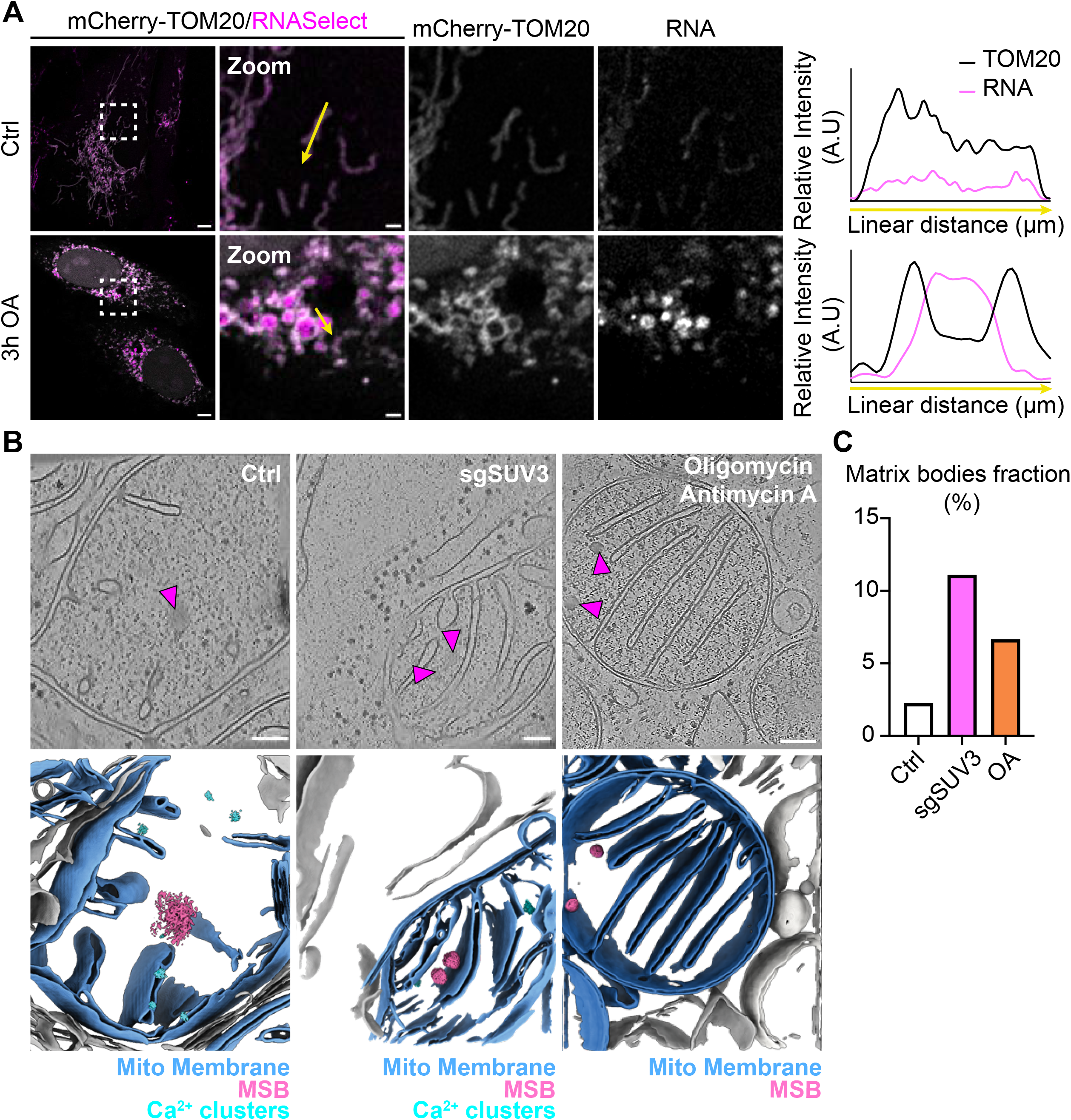
General mitochondrial stress through ETC inhibition is associated with stress body formation in a subset of mitochondria. **(A)** Live imaging of control and OA-treated U2OS cells transfected with mCherry-TOM20 and stained with RNASelect. OA treatment was performed for 3 hours with a mixture of 3uM oligomycin and 3uM antimycin A. Scale bars: 5 μm; 1 μm in zoom. A single slice of a z-stack is shown for each condition. Right: Relative mCherry-TOM20 and RNASelect intensity plots for the yellow arrows drawn on the left. **(B)** Cryo-ET tomograms of control, sgSUV3, and OA-treated cells with mitochondrial bodies. Scale bar: 100nm. OA-treated images were previously acquired in *Rose et al.*, 2025 (32). **(C)** Quantification of matrix bodies fraction occupied for each condition in (B). The number of mitochondria with aggregates compared to the total number of mitochondria analyzed for each condition is 1/44 for control, 4/36 for sgSUV3, and 3/45 for OA.

Using *in situ* cryo-ET, we recently observed the nanostructural consequences of OA-induced mitochondrial dysfunction on mitochondrial ultrastructure in cells expressing Parkin (32). As in SUV3 depletion, OA treatment was associated with reorganization of the IMM and decreased cristae density. Given our live-imaging results, and the similar impacts of OA and SUV3 depletion on mitochondrial structure, we then considered the idea that general mitochondrial dysfunction is also associated with the presence of protein aggregates. We reanalyzed our previously acquired *in situ* cryo-ET tomograms, and found the presence of mitochondrial aggregates in a subset of the mitochondria analyzed suggesting that protein aggregates (**Fig. 3B-C**) (32). This suggests that protein aggregates might be a common feature of mitochondrial perturbations linked to mitochondrial RNA organization. Collectively, these findings reveal that mitochondrial ultrastructural remodeling concurrent with MSB formation is a common, generalizable feature of mitochondrial dysfunction.

### Mislocalization of ATP synthase to the inner boundary membrane defines cristae remodeling in SUV3-depleted cells

In homeostasis, the dimerization of mitochondrial ATP synthase contributes to IMM curvature required to maintain cristae morphology, characterized by deep invaginations of the IMM, that house the electron transport chain complexes (40–42). In addition to the cristae proper, the IMM is also composed of the inner boundary membrane (IBM), a flat subregion that is typically adjacent to the OMM, and the cristae junctions (CJ) that connect IBM to cristae and act as diffusion barriers (41). We previously found that, in cells treated with mitochondrial poisons oligomycin and antimycin A to prompt mitochondrial depolarization, ATP synthase molecules are mislocalized from the cristae to the IBM, consistent with loss of IMM organization and subdomain identity (32).

To ask whether mislocalization of ATP synthase could underlie the ultrastructural consequences of SUV3-depletion, we visualized endogenous ATP Synthase subunit ATP5B using indirect immunofluorescence labeling. We compared TOM20, dsRNA, and ATP5B signals between fixed control and SUV3-depleted U2OS-Cas9 cells, revealing heterogeneous localization of both dsRNA and ATP synthase among SUV3-depleted mitochondria (**Fig. 4A**). In control cells, dsRNA levels were low-to-undetectable, and ATP5B was homogeneously distributed along individual mitochondria (**Fig. 4A-B**). Mitochondria containing “straight” cristae were the minority in SUV3-depleted cells (**Fig S4C-D**). In contrast, SUV3-depleted mitochondria containing abundant dsRNA could be divided into subsets in which dsRNA concentrated in a dense core in the matrix, or in which dsRNA collapsed parallel to the OMM respectively; we refer to these as “sectored” and “collapsed” mitochondria (**Fig. 4A-B and Fig. S4A-B**).

**Figure 4:**
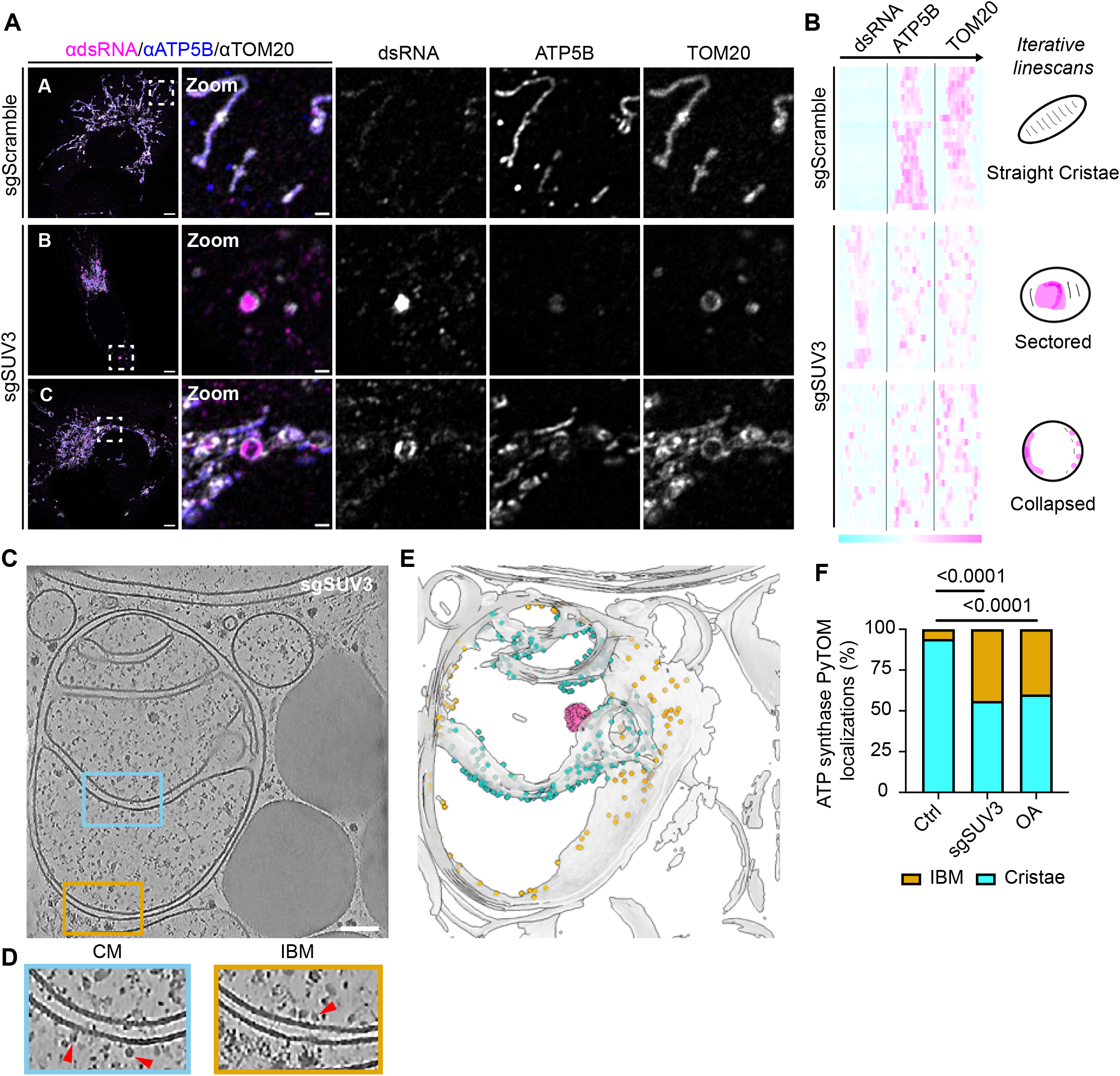
dsRNA and ATP5B are differently organized across the mitochondrial network following sgSUV3 stress. **(A)** Immunofluorescence images of U2OS-Cas9 transfected with sgScramble (top) and sgSUV3 (bottom), and immunolabeled with anti-TOM20, anti-dsRNA (J2), and anti-ATP5B. Imaged 7 days after transfection. The top row represents an example of straight mitochondria (A), the middle row represents an example of sectored mitochondria (B), and the bottom row represents an example of collapsed mitochondria. Scale bars: 5 μm; 1 μm in zoom. A single slice of a z-stack is shown for each condition. **(B)** Relative intensity heatmaps of TOM20, dsRNA, and ATP5B for mitochondrial phenotypes represented on the right. The heatmaps are derived from linescan analysis of the experiment in A. Each row represents a linescan with the represented heatmap intensities scaled to each linescan. 63 mitochondria were analyzed across three biological replicates, with 21 mitochondria per category and 7 mitochondria per replicate for each category. **(C)** Cryo-ET image of U2OS cells transfected with sgSUV3. Scale bar: 100nm **(D)** Insets shown in (C) showing examples of ATP synthase molecules localized to the cristae membrane (CM) or the inner boundary membrane (IBM). Red arrows point to ATP5B dimers. **(E)** 3D segmentation of the cryo-ET tomogram shown in (C). Blue dots represent ATP synthase localized to the CM, and orange dots represent ATP synthase mislocalized to the IBM. A matrix body is shown in magenta. **(F)** Quantification of high correlation PyTOM localization of control, sgSUV3 and oligomycin/antimycin A (OA) treated U2OS cells. Cryo-ET images of sgSUV3 cells were acquired in experiment from panel (C) and OA treated and control samples were reanalyzed from tomograms previously acquired in *Rose et al.*, 2025 (32). The analysis was performed on a single representative tomogram containing hundreds of individual ATP synthase molecules. N = 17 IBM and 623 cristae for control, N = 179 IBM and 232 cristae for sgSUV3, N = 257 IBM and 413 cristae for OA. Statistical significance was determined using Fisher’s Exact test.

To analyze the relationship between dsRNA and ATP synthase localization, we measured the position of dsRNA fluorescence maxima relative to ATP5B or TOM20 signals, representing profiles of each marker within individual mitochondria with an intensity heatmap (n=84). In control cells, ATP5B signals were most intense along the central axis of each mitochondrion, consistent with the known localization to the highly curved cristae membrane (**Fig. 4B**, upper row). However, ATP5B signal collapsed to the organelle periphery in SUV3-depleted cells; this was particularly prominent for the subset of mitochondria in SUV3 knockouts that lacked MSB (**Fig. 4B**, middle and lower rows, **Fig. S4A-B**).

These data suggested that ATP synthase is mislocalized in SUV3-depleted cells and that this is correlated with cristae collapse independent of local dsRNA abundance. To test this, we used *in situ* cryo-ET of SUV3-depleted cells and quantified the number of ATP synthase molecules in the cristae membrane and IBM within tomograms by performing 3D template matching of ATP synthases using a reference generated from our data (*Rose et al., 2025*) (**Fig. 4C)** (32). We observed a significant relocalization of ATP synthase molecules in SUV3-depleted mitochondria from the cristae membrane toward the IBM (**Fig. 4D-E)**.

Quantification of the PyTOM localization and comparison to our previously published dataset on ATP synthase mislocalization during mitochondrial depolarization by OA revealed that IMM organization in SUV3-depleted cells structurally mimics the loss of membrane integrity during depolarization (**Fig. 4F**) (32). Together, these findings suggest that ATP synthase mislocalization underlies cristae collapse and loss in a subset of mitochondria.

### Spatial compartmentalization of oxidative and reductive components within mitochondrial networks

Given our observation of ATP synthase mislocalization in SUV3-depleted cells, we next considered how metabolic remodeling previously linked to cristae loss could relate to mitochondrial stress body formation and/or localization in cells. Thus far, we have shown that SUV3 depletion impacts nucleic acid and protein components linked to the oxidative functions of mitochondria, including mitochondrial RNA species, mitoribosomal protein MRPL23, and ATP synthase. However, mitochondria also participate in the metabolic pathways of reductive synthesis (13).

Pyrroline-5-carboxylate synthase (P5CS), the rate-limiting enzyme in reductive proline and ornithine synthesis, is at the intersection between oxidative and reductive clusters and can oligomerize to form filaments that are sequestered within a subset of mitochondria during periods of elevated bioenergetic demand (**Fig. 5A**) (18). Interestingly, increasing OXPHOS demand exacerbates the P5CS filamentous morphology and specifically leads to sequestration of the enzyme within mitochondria that lack ATP synthase and cristae in a manner dependent on mitochondria fusion-fission dynamics (18). Given our observation of cristaeless mitochondria that lacked MSB within SUV3-depleted cells, we next considered a model in which MSB sequester oxidative mitochondrial components for sorting into a subset of mitochondria distinct from the reductive P5CS filaments (**Fig. 5A**).

**Figure 5:**
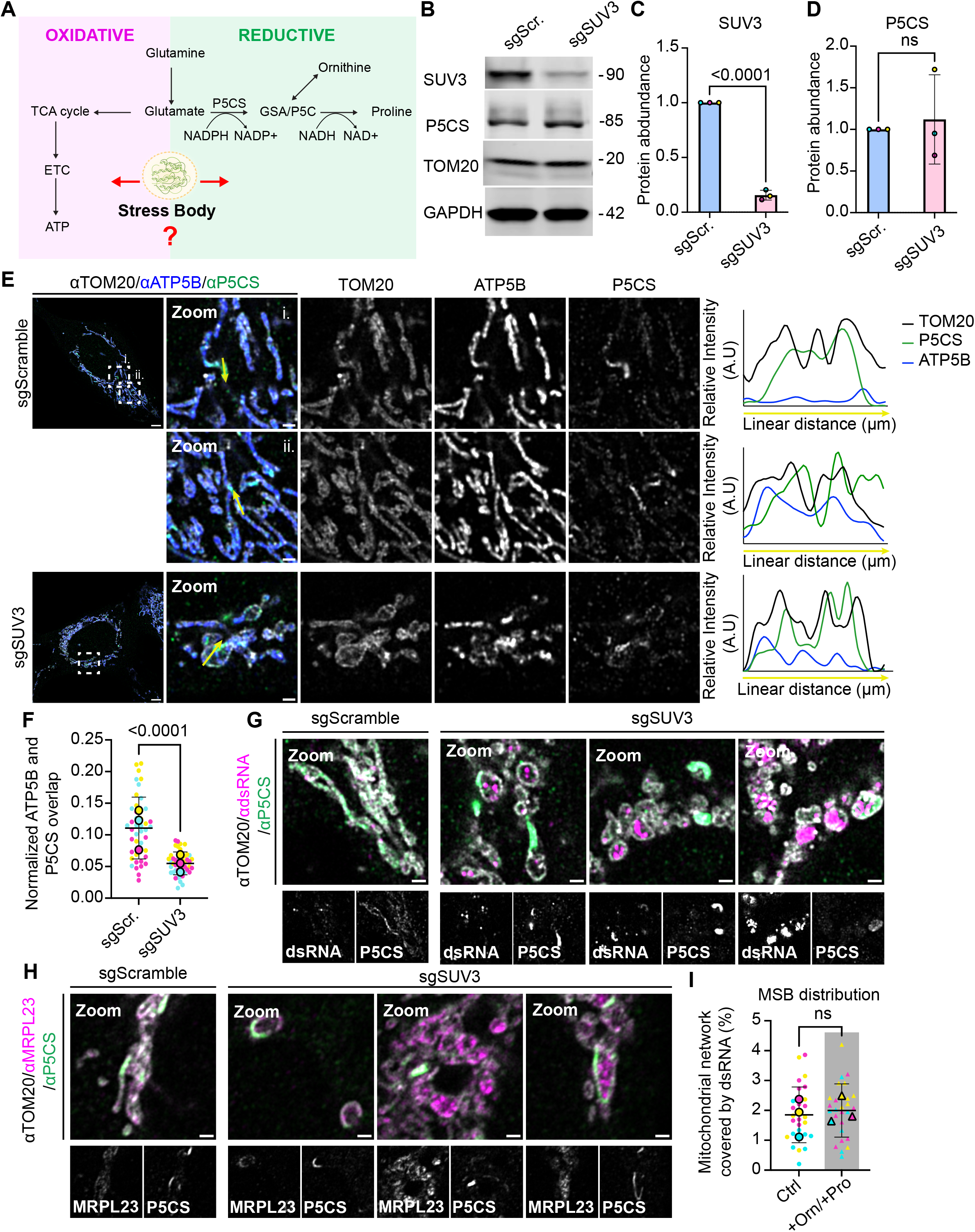
P5CS filaments and MSBs appear in distinct mitochondrial subdomains. **(A)** Representation of oxidative and reductive mitochondria processes. **(B)** P5CS, TOM20, and GAPDH protein levels in sgScramble versus sgSUV3 U2OS-Cas9 cells analyzed by western blot. GAPDH as the loading control for three independent biological replicates. **(C)** Quantification of the experiment performed in (B) for SUV3 abundance. **(D)** Quantification of the experiment performed in (B) for P5CS abundance. For (C) and (D), the protein abundance is normalized to TOM20 and shown as a fold change relative to sgScramble. N = 3 independent biological replicates. Error bars display mean values ± standard deviation. Statistical significance determined with an unpaired t-test. **(E)** Left: Immunofluorescence images of U2OS-Cas9 transfected with sgScramble (top) and sgSUV3 (bottom), and immunolabeled with anti-TOM20, anti-ATP5B, and anti-P5CS. Imaged 7 days after transfection. Right: Relative TOM20, ATP5B, and P5CS intensity plots for the yellow arrows drawn on the left. Scale bars: 5 μm; 1 μm in zoom. A single slice of a z-stack is shown for each condition. **(F)** Quantification of the experiment performed in (E) showing the Jaccard Index, labeled as overlap. Quantification performed on maximum intensity projection images. N = 42 FOV for sgScramble and N = 46 FOV for sgSUV3 over three independent replicates. Error bars display mean values ± standard deviation, and the bottom bar is clipped if below 0. Statistical analysis performed on individual data points with Mann-Whitney test. **(G)** Immunofluorescence images of U2OS-Cas9 transfected with sgScramble (i.) and sgSUV3 (ii., iii., iv.), and immunolabeled with anti-TOM20, anti-dsRNA (J2), and anti-P5CS imaged 7 days after transfection. Scale bars: 5 μm; 1 μm in zoom. A single slice of a z-stack is shown for each condition. **(H)** Immunofluorescence images of U2OS-Cas9 transfected with sgScramble (i.) and sgSUV3 (ii., iii., iv.), and immunolabeled with anti-TOM20, anti-MRPL23, and anti-P5CS imaged 7 days after transfection. Scale bars: 5 μm; 1 μm in zoom. A single slice of a z-stack is shown for each condition. **(I)** Quantification of total dsRNA area covering the mitochondrial network in control sgSUV3 cells and sgSUV3 cells with excess ornithine and proline. Quantification performed on maximum intensity projection images from the experiment in Fig. S8B. N = 27 FOV (control), 26 FOV (+Orn/+Pro) across three independent biological replicates. Error bars display mean values ± standard deviation, and bottom bar is clipped if below 0. Statistical analysis performed with Mann-Whitney test on individual data points.

After validating that P5CS protein abundance is not dependent on SUV3 function (**Fig. 5B-D; Fig. S5-6**), we considered the localization of P5CS filaments relative to either ATP5B or dsRNA within mitochondrial networks in control versus SUV3-depleted U2OS cells immunolabeled with antibodies against ATP5B, P5CS, and TOM20. Colocalization between ATP5B and P5CS decreased in SUV3-depleted cells (**Fig. 5E-F**), consistent with P5CS clustering in mitochondrial domains lacking ATP synthase. P5CS signals and MSB components were sequestered into distinct mitochondrial subdomains at both the organelle and mitochondrial network levels (**Fig. 5G-H; see uncropped images in, S7**). We noted the presence of P5CS filaments that collapsed parallel to the OMM marker TOM20 in a subset of mitochondria, while dsRNA and MRPL23 could still be observed within the matrix (**Fig. 5G-H, S7**). Overall, these findings are consistent with spatial compartmentalization of reductive and oxidative markers across the mitochondrial network during mitochondrial RNA processing stress.

### Mitochondrial stress bodies and P5CS filaments are independently regulated following stress compartmentalization

The assembly of P5CS into filament structures is thought to enhance the efficiency of its enzymatic activity in the reductive synthesis of ornithine and proline (18,43,44). Cell culture in the presence of excess ornithine and proline causes the dissolution of P5C oligomers and P5CS monomer diffusion throughout the mitochondrial network, consistent with the idea that filament persistence is controlled in part by substrate channeling and the output of reductive synthesis (**Fig. S8A**) (18). Given this visual marker of P5CS regulation, we sought to test whether the subset of mitochondria containing oxidative versus reductive components may be differentially regulated. We supplemented either control or SUV3-depleted cells with excess ornithine and proline then asked whether P5CS filament dissolution led to concurrent remodeling of dsRNA-positive MSB in the matrix. In both control and SUV3-depleted cells, ornithine and proline supplementation shifted P5CS distribution from the filamentous pattern to a diffuse localization as expected (**Fig. S8B-C**). We then quantified mitochondrial dsRNA to determine whether ornithine and proline supplementation led to redistribution of dsRNA, and thus MSB. Consistent with independent regulation of oxidative and reductive components, ornithine and proline supplementation had no effect on the fraction of mitochondrial network occupied by dsRNA in SUV3-depleted cells (**Fig. 5I**). To validate that modulation of reductive pathway does not affect dsRNA distribution, we supplemented SUV3-depleted cells with glutamine in a distinct parallel experiment, which may be typically utilized by P5CS for ornithine and proline synthesis, and again assessed dsRNA distribution. Consistent with our previous observations, glutamine supplementation did not influence dsRNA distribution (**Fig. S9**). Together, our findings indicate that during mitochondrial RNA processing stress, the sequestration of gene expression components within MSB is a distinct and independently regulated post-transcriptional response, that co-exists with the P5CS oligomerization response to bioenergetic demand but in two distinct subsets of mitochondria within the cell.

Collectively, these data support a model in which SUV3-depletion results in the formation of mitochondrial stress bodies that contain components linked to oxidative functions within the cristae-rich domains of the mitochondrial network, while within the same cell, the reductive mitochondrial compartments marked by oligomerized P5CS are independently regulated in cristae-depleted domains devoid of ATP synthase (**Fig. 6**).

**Figure 6:**
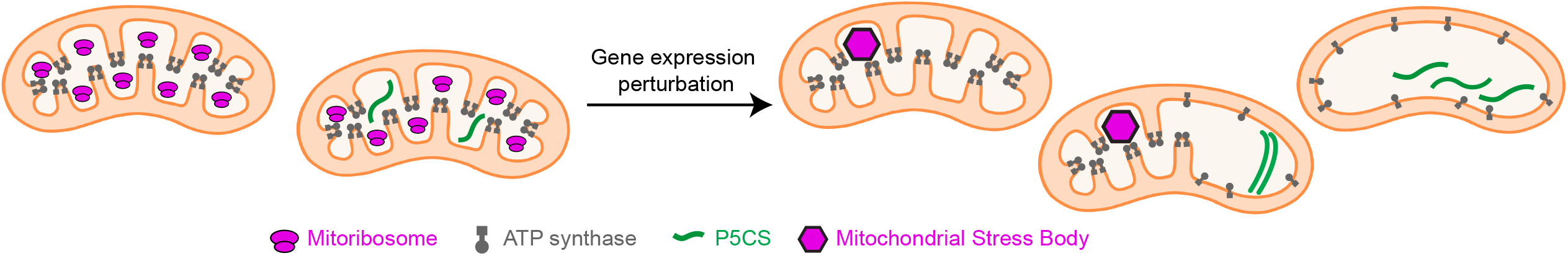
MSB formation in oxidative mitochondrial domains following sgSUV3 stress. Schematic illustrating the spatial compartmentalization of oxidative and reductive components under sgSUV3 stress.

## DISCUSSION

Mitochondrial heterogeneity is omnipresent within the mitochondrial network (45,46). In a eukaryotic cell, this network can comprise up to thousands of mitochondria that can differ by their shape, content, functional state, and fates (45–48). Our data indicate that RNA processing stress is associated with a heterogeneous reorganization of mitochondrial proteins and RNAs across the mitochondrial network that is accompanied by restructuring of the IMM. In this study, we applied *in situ* cryo-electron tomography to gain a nanoscale view of mitochondrial inner membrane remodeling that facilitates MSB formation when cells are in crisis. This allowed the first-ever high-resolution examination of mitochondrial stress bodies and their relationship to the mitochondrial inner membrane *in situ*.

ATP synthase mislocalization was a significant feature of the ultrastructural changes observed in SUV3-depleted cells. Strikingly, we observed inner membrane domains that contained MSB within intact cristae, while mitochondria that lacked MSB had vesicular, collapsed, or looping cristae. These observations suggest at least two testable models: either (1) MSB may serve a protective role by sequestering mitochondrial RNA, misfolded proteins, and disassembled mitoribosome components like MRPL23 away from the rest of the proteome of the mitochondrial network; or (2) cristaeless mitochondria exclude stress bodies in favor of other structures, namely P5CS oligomers, which are known to congregate within a subset of mitochondria devoted to the reductive synthesis of ornithine and proline.

Our data are consistent with a model in which RNA processing stress promotes the coalescence of components associated with oxidative function into MSBs that localize to domains that are spatially distinct from reductive bodies. These findings suggest the possibility that the codification of mitochondria into distinct classes could inform the fate of individual organelles during stress management and recovery, if for example, MSB-positive or P5CS filament-positive mitochondrial subsets become preferential targets for autophagy during stress.

Here, we observed that OA treatment is associated with protein aggregate presence as well as with mitochondrial RNA reorganization in a subset of swollen mitochondria. This indicates that MSB features can be shared by distinct mitochondrial stressors, and it is important to further characterize how these bodies are similar or dissimilar between distinct conditions. Mitochondrial dysfunction is common to a broad variety of diseases (49). Understanding how distinct stressors impact mitochondrial organization will be critical to strengthen our comprehension of mitochondrial homeostasis, and to refine treatment strategies in view of similarity or dissimilarities between stressors.

We found that P5CS oligomers/filaments reside in mitochondrial compartments that are distinct from those that contain MSB, suggesting that mitochondrial components associated with oxidative functions are sequestered within a separate mitochondrial network compartment from the reductive components during stress. We further show that P5CS oligomerization is regulated independently of MSB coalescence, indicating that MSB and P5CS oligomers serve parallel functions during mitochondrial stress: protecting the mitochondrial gene expression machinery in the case of MSB, and tuning the efficiency of reductive synthesis in the case of P5CS structures. The separation of P5CS in the sub-network domains is known to require cycles of fusion and fission (18). Thus, mitochondrial dynamics are likely to play a role in the seeding of MSBs in the early stages of stress, and for compartmentalization of oxidative stress away from reductive components.

Our data further indicate that cristae disorganization is heterogeneous across the network, and points towards a potential protective role of MSBs in maintaining cristae ultrastructure. Previous work linked a mitochondrial genome mutation to nascent chain proteotoxicity in the matrix, though the mechanism of mitochondrial RNA surveillance that linked the two was unclear (50). Our data suggest that mitochondrial stress body formation downstream of SUV3 loss-of-function may be a means to preserve local IMM ultrastructure, and potentially, ETC function (9). Mitochondrial protein aggregates induced by inhibition of HSP90 suggested that, in the context of proteotoxic stress, aggregates drive inner mitochondrial membrane cristae collapse (29). Our data suggest the opposite, which raises the possibility that there are multiple types of bodies/aggregates within the mitochondrial matrix that serve distinct roles or act in a context-dependent manner. Mitoribosomal proteins are particularly prone to become insoluble in the matrix during HSP90 inhibition, especially when the mitochondrial UPR is impacted (33). It would be interesting to determine whether this aggregation of proteins is accompanied by mitochondrial RNA reorganization, and to identify other mitochondrial stressors associated with both RNA and protein reorganization within the matrix.

Future work to manipulate matrix aggregates or P5CS oligomers such as by synthetically inducing the formation of mitochondrial aggregates or condensates would allow direct assessment of their impact on cristae organization and mitochondrial proteome quality control during a variety of stress conditions; directly assessing the RNA content of aggregates induced by proteotoxic stress; identifying the complete set of proteins found within MSB using biochemical and proteomic methods; and determining the genes and pathways essential to MSB resolution after stress abatement.

Further *in situ* examination of mitochondrial cristae integrity may be particularly fruitful, as these membranes preserve the homeostatic mitochondrial localization of nucleic acids that could otherwise stimulate inflammatory responses (6,51). While our tomography data did not reveal obvious formation of mitochondrial membrane pores such as those mediated by VDAC or BAK/BAX, further assessment of the potential cytosolic release of mitochondrial nucleic acids will likely be informative if, for example, sequestration of mitochondrial nucleic acid in MSB may in fact restrict its release to the cytosol through damaged or permeabilized IMM which would integrate MSBs formation and potentially disassembly with cytosolic innate immune surveillance pathways (52–54).

Overall, the main advance of our study is to demonstrate the coexistence of multiple classes of distinctly composed and remodeled mitochondria within the same cytoplasm. Interestingly, we observed a third set of mitochondria that contained neither MSB nor P5CS oligomers/filaments. This third set may comprise mitochondria that are in the process of remodeling into either of the other two types, or it could represent another class of mitochondrion that has a yet-unknown function within the mitochondrial consortium.

## SUPPLEMENTAL FIGURES

Figure S1: Mitochondrial organization of MRPL23 and dsRNA within stress bodies.

Figure S2: Process of acquiring cryo-ET tomograms.

Figure S3: SUV3 depletion induces inner mitochondrial membrane remodeling in live cells.

Figure S4: Mitochondrial phenotype variety in sgSUV3 cells.

Figure S5: Protein levels of mitochondrial proteins of interest following SUV3 depletion.

Figure S6: Uncropped western blots.

Figure S7: Localization of P5CS and MSB components following SUV3 depletion.

Figure S8: Ornithine/Proline supplementation does not affect dsRNA distribution.

Figure S9: Glutamine supplementation does not impact dsRNA distribution.

## SUPPLEMENTAL TABLES

### Table S1: Key Resource Table

Table S2: Sequences of guide RNAs for the generation of CRISPR knockouts.

Table S3: Sequences of primers used for CRISPR knockout validation.

Table S4: CRISPR KO efficiency across seven experimental replicates.

## METHODS

### Mammalian cell growth and amino acid supplementation

Human U2OS cells (RRID:CVCL_0042) and U2OS-Cas9 cells (Horizon, HD Cas9-012) were received from the UC Berkeley Cell Culture facility. Cells were cultured for no more than 15 passages in high-glucose, no glutamine Dulbecco’s modified Eagle’s medium (DMEM - Thermo Fisher Scientific, #11960044) supplemented with 10% heat-inactivated fetal bovine serum (Thermo Fisher Scientific, #A3840101), 1% penicillin/streptomycin (Gibco, Thermo Fisher Scientific, #15140122), and 2mM L-glutamine (Thermo Fisher Scientific, #25030081) at 37°C in a humidified 5% CO_2_ chamber. For experiments in which cells were treated with exogenous amino acids, heat-inactivated fetal bovine serum was replaced by dialyzed FBS (Thermo Fisher Scientific, 26400044) and L-glutamine was replaced by 2mM or 10mM of GlutaMax (Gibco, Thermo Fisher Scientific, 35050061). Treatment with proline and ornithine was done for 4 hours before fixation by adding 5mM of L-proline (Sigma-Aldrich, P0380) and 5mM of L-ornithine monohydrochloride (Sigma-Aldrich, O2375) to the DMEM media.

### Generation and validation of CRISPR knockouts

Guide RNA sequences used for non-targeting controls and to target SUPV3L1 are reported in table S2. Knock-out cells were generated using a modified version of the Lipofectamine CRISPRMAX Cas9 Transfection Reagent protocol (Thermo Fisher Scientific, CMAX00008). Lipofectamine transfection was done in 6-well plates once cells reached 30%-60% confluency. For CRISPR editing of U2OS cells (used in the *in situ* cryo-ET experiments), the transfection mixture consisted of 2uL of 20uM NLS-Cas9 (Synthego) mixed with 5.2uL of 10uM sgRNAs (Gene Knockout Kit, EditCo) and 12.8uL of Opti-MEM (Thermo Fisher Scientific, REF: 31985-070) for a final volume of 20uL of Cas9/sgRNA solution. Parallelly, 10uL of Cas9 Plus reagent was mixed with 70uL of Opti-MEM and then added to the tube with the complexed Cas9/sgRNA. Finally, 12uL of Lipofectamine CRISPRMAX Cas9 reagent was diluted with 88uL of Opti-MEM, mixed, and added to the Cas9/sgRNA/Cas9 Plus mixture. The final mixture was added dropwise to the plated cells following a 10 minutes incubation period at room temperature. For CRISPR editing of U2OS cells stably expressing Cas9 (U2OS-Cas9), the transfection mixture was prepared by mixing 4uL of 10uM synthetic guide RNAs (Gene Knockout Kit, EditCo) with 150uL of Opti-MEM. In parallel, 9 μL of Lipofectamine RNAiMax (Thermo Fisher Scientific, REF: 13778075) was mixed with 150 μL of Opti-MEM in another tube, and 150uL of this mixture was subsequently added to the synthetic guide RNA transfection mixture. The mixture was incubated for 5 minutes at room temperature, and 250uL of the mixture was then added in dropwise-manner to the plated cells. For both U2OS and U2OS-Cas9 cells, the media was replaced with fresh media 24 hours after the transfection, and experiments or fixation were performed 7 days post-transfection. Protein depletion following transfection with the synthetic guides RNA was validated using western blot in three independent biological replicates. Genome editing efficiency was estimated by Sanger sequencing in seven of the experimental replicates, including three that were analyzed by western blot. To prepare sequencing samples, phenol-extracted genomic DNA was PCR-amplified using Phusion High-Fidelity DNA Polymerase (New England Biolabs) with the forward and reverse PCR primers (IDT) listed in table S3. The PCR products were then resolved on a 1.5% agarose gel containing SYBR Safe DNA stain (Thermo Fisher Scientific), purified with the commercial kit ZYMO DNA Clean & Concentrator-5 (Zymo; D4004) and submitted to the UC Berkeley DNA Sequencing Facility for BigDye Sanger sequencing with forward or reverse nested sequencing primers (IDT) listed in table S3. Publicly available ICE CRISPR Analysis. 2025. v3.0. EditCo Bio was used to confirm the presence of indels and estimate knockout efficiency (table S4).

### Western Blot

Cells were first trypsinized and centrifuged at 800 x g for 5 minutes. The supernatant was then carefully removed, and cells were gently resuspended with PBS, and centrifuged again at 800 x g for 2 minutes. The cells were then lysed using lysis buffer (50mM Tris-HCl, pH 7.4, 150mM NaCl, 1mM EDTA, 1% Triton X-100, and ultrapure distilled water) supplemented with a protease inhibitor cocktail (Roche, 11697498001) and kept on ice for 10 minutes. Then, the cells were centrifuged at 13,000 x g for 10 minutes, and the supernatant containing the proteins was collected to perform protein concentration estimation using DC protein assay (BioRad). For SDS-PAGE, β-mercaptoethanol (BME) was added to 2× Laemmli sample buffer at a final concentration of 10% (v/v; 5 µL BME per 45 µL Laemmli buffer). Protein samples were then mixed with the prepared Laemmli/BME buffer at a 1:1 (v/v) ratio and heated at 90°C for 5 min. The solution was subjected to SDS-PAGE for electrophoresis using precast gels (Invitrogen Cat. No. XP04205BOX), and the proteins were then transferred to a nitrocellulose membrane using the iBlot NC mini-stack (Invitrogen Cat. No. IB23002). The SDS-PAGE gel was stained overnight with Coomassie Brilliant Blue at 4°C, washed with ultrapure distilled water, and used as an additional qualitative check of protein distribution across lanes. The nitrocellulose membrane was placed in blocking buffer and blotted overnight at 4°C with the following antibodies (1:5000): rabbit SUPV3L1 Polyclonal antibody (Proteintech, #12826-1-AP), mouse anti-MT-ATP6 (Abcam, #ab219825), mouse anti-P5CS Monoclonal antibody (Proteintech, 68184-1-Ig), mouse anti-mt OPA1 (Novus Biologicals, NBP171656), rabbit anti-TOMM20 (Abcam #ab186735), mouse anti-mtHSP70 Monoclonal Antibody (JG1) (Invitrogen #MA3-028), rabbit anti-MRPL43 (Invitrogen, # PA5-103527), mouse anti-TFAM (Abcam, #ab307302), rabbit anti-Mitofilin (Abcam, #ab137057), and mouse anti-GAPDH (Sigma-Aldrich, CB1001-500UG). After the primary incubation, the membrane was washed with TBST and incubated for 1 h at room temperature with the secondary antibodies (1:5000) diluted in blocking buffer: goat anti-rabbit IgG (H+L), IRDye 680RD conjugate (LI-COR, Cat. No. 926-68171), and goat anti-mouse IgG (H+L), Alexa Fluor Plus 800 conjugate (Invitrogen, Cat. No. A32730). After the secondary incubation, the membrane was washed with TBST and visualized using Licor Odyssey Clx Imager.

### Fixation, antibodies and immunofluorescence

Prior to fixation, cells were seeded onto glass-bottom 35-mm dishes (Mattek) and cultured for 24 hours. Cell fixation was performed using pre-warmed (37 °C) 4% paraformaldehyde solution made in pH 7.4 Phosphate Buffered Saline (PBS) for 30 minutes at room temperature and protected from light. The cells were then washed with PBS prior to permeabilization in 0.1% Triton X-100 diluted in PBS for 10 minutes. Glass-bottom dishes were then washed with TBST (tris-buffered saline with 0.1% Tween 20) blocking buffer (pH 7.5) containing 1% bovine serum albumin for 10 minutes or 1 hour (for samples that would be stained with ATP5B-488) on an orbital shaker. The TBST buffer was then replaced by a TBST buffer containing the primary antibodies at the dilution ratios specified below and dishes were incubated overnight at 4°C. The dishes were then washed with TBST buffer for 10 minutes on an orbital shaker prior to a 1hr room temperature incubation with TBST buffer containing secondary antibodies at the concentrations specified below. We detected endogenous proteins and RNA using the following antibodies: rabbit anti-MRPL23 at 1:1000 (Sigma-Aldrich, HPA050406), mouse anti-J2 at 1:1000 (Sigma-Aldrich, MABE1134), rat anti-TOMM20 at 1:500-1:1000 (Abcam, ab289670), mouse anti-ATP5B at 1:1000 (Invitrogen, A-21351), rabbit anti-P5CS at 1:500 (Proteintech, 17719-1-AP), mouse anti-P5CS at 1:1000 (Proteintech, 68184-1-Ig), rabbit 488-conjugated ATPB at 1:1000 (Proteintech, CL488-17247), goat anti-Rat IgG (H+L) Highly Cross-Adsorbed Secondary Antibody Alexa Fluor™ Plus 405 (Thermo Fisher Scientific, A48261), Goat anti-Rat IgG (H+L) Highly Cross-Adsorbed Secondary Antibody, Alexa Fluor™ Plus 488 (Thermo Fisher Scientific, A48262TR), Donkey anti-Rabbit IgG (H+L) Highly Cross-Adsorbed Secondary Antibody, Alexa Fluor™ Plus 680 (Thermo Fisher Scientific, A32802), Donkey anti-Rabbit IgG (H+L) Highly Cross-Adsorbed Secondary Antibody, Alexa Fluor™ Plus 488 (Thermo Fisher Scientific, A32790TR), Goat anti-Mouse IgG (H+L) Highly Cross-Adsorbed Secondary Antibody, Alexa Fluor™ 568 (Thermo Fisher Scientific, A-11031). More details on the immunolabeling protocol can be found here: https://dx.doi.org/10.17504/protocols.io.n92ldr1xxg5b/v1

### Transient transfection, OA-treament, and vital dyes

Transient transfection was performed as previously described (https://dx.doi.org/10.17504/protocols.io.ewov1dr82vr2/v1) with the modification that cells were plated onto the imaging dishes on the day of transfection instead of the day prior to transfection. Transient transfection was performed using Opti-MEM™ I Reduced Serum Medium (Thermo Fisher Scientific, REF: 31985-070), Lipofectamine 2000 reagent (Thermo Fisher Scientific, REF:11668030), and 500ng of mCherry-TOMM20-N-10 obtained from Addgene (Addgene plasmid # 55146). On the day of imaging, the media was replaced with either fresh media or media containing a 3uM:3uM mixture of Oligomycin (Sigma catalog: SIAL-O4876-5MG) and Antimycin A (Sigma catalog: SIAL-A8674-25MG) for 3 hours. After 2h20 of incubation, 1mL of media was saved in a separate tube while 5uM of RNA Select (Invitrogen, S32703) was added to the plated cells in the remaining 1mL of media. The cells were incubated with RNA select for 30 minutes, and then the media was replaced with the saved media, and imaging was performed at the 3-hour timepoint. A detailed protocol might be found here: https://dx.doi.org/10.17504/protocols.io.r3l25my1jl1y/v1. For live cell-imaging of MitoTrackerGreen, 1mL of conditioned media was removed and saved and then 200nM of MitoTracker Green (Invitrogen, M7514) was directly added to the dish for 30 minutes and then replaced with the conditioned media before imaging. For *in situ* cryo-ET, the U2OS cells were fluorescently labeled with 25nm of MitoTracker Deep Red (Invitrogen, M22426) and 5uM of RNA Select for 30 minutes, followed by a media wash. The cells were then prepared for *in-situ* cryo ET as described below, and fluorescence labeling did not direct data acquisition.

### Airyscan fluorescence microscopy and image acquisition

Confocal imaging was performed using the Zeiss LSM 980 with Airyscan 2 laser scanning microscope, equipped with the laser lines 405, 488, 561, and 639 nm. Image acquisition was performed using an inverted 63x/1.4 numerical aperture oil objective. Live-cell imaging was conducted in a humidified chamber maintained at 37°C with 5% CO_2_. 3D Airyscan processing and maximum projections were generated using the Zeiss ZEN Blue software version 3.7 (Carl Zeiss). Zen Blue and FIJI were used to linearly adjust brightness and/or contrast.

### Image analysis

Arivis Vision4D (ver 4.2.1) (RRID:SCR_018000) was used for the analysis of maximum projection czi files after sis format conversion, while FIJI was used for the line scan analysis of individual slices of 3D airyscan processed czi files.

#### Per cell analysis

Images containing a single cell were directly analyzed. For images containing more than one cell, the draw object tool of Arivis was used to individually segment cells as objects. An object mask was then applied, and the output was saved as a separate image. In the case of two cells, the cells were either both individually segmented prior to the object mask or an invert mask was used following segmentation of the first cell.

#### Segmentation

Segmentation of TOM20 and TOM20-enclosed segments, defined as the mitochondrial network, was performed using Arivis’s intensity threshold segmenter with a manual threshold. Internal holes were preserved for the TOM20 segmenter to allow the mitochondrial inclusion of structures of interest in subsequent steps. Mitochondrial segments were then filtered based on area with a cutoff of 400,000 nm^2^. MRPL23, ATP5B, and J2 were segmented using intensity threshold segmentation, with internal holes allowed, and without additional size filtering steps. Thresholds were defined with Otsu’s automatic method for MRPL23, and manually for ATP5B, P5CS, and J2. When intensity variation across replicates prevented the use of a fixed threshold, J2 was segmented using Otsu. All objects were then filtered based on mitochondrial localization using Arivis compartmentalization function with a cutoff of 50%, and objects situated within the TOM20-enclosed segments were defined as “mitochondrial objects”. Following segmentation, information such as area measurements were exported as csv files. Images in which TOM20-segmentation resulted in inclusion of non-mitochondrial regions were excluded. When Otsu was used for J2 segmentation, the mean segmented J2 intensity across all sgScramble images of each replicate was calculated. sgSUV3 images were excluded if their mean J2 intensity was lower than the mean segmented J2 intensity of the corresponding sgScramble control replicate. For quantification of the confocal microscopy data, data points that were over or under 3SD from the mean were excluded.

#### Object overlap measurements

To calculate overlap between objects, the object math function of Arivis was used and the area measurements of the overlapping regions and separate structures were saved. Then, Jaccard Index was measured by the following equation:

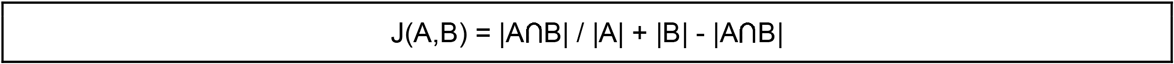

With A and B corresponding to the area of the structures of interest and the intersection of A and B corresponding to the overlapping region. To determine the proportion of mitochondrial network covered by a mitochondrial object of interest, the total area of the mitochondrial object of interest was divided by the total area of the mitochondrial network. The values were then computed as a proportion of the mitochondrial network occupied in %. Channels exhibiting detector saturation were retained for object segmentation and overlap analyses.

#### Line scan analysis

Line scans were performed on individual slices of Airyscan-processed images using FIJI. Straight lines were drawn using the line tool and saved in the ROI manager. Intensity profiles for each channel were obtained using the Plot Profile function, and the plot values were exported as csv. The data was then imported into GraphPad prism (version 11.0.0) for representation or organized and saved in tab delimited files for import in R and generation of heatmaps. In images where filaments were partially saturated, line scans were restricted to filaments in which the signal remained within the detector’s dynamic range. Linescans could be of different length except for the heatmap generation where all linescans were of the length of 3.34 µm. Linescans were drawn over the minor and major axis of the mitochondria.

### Cryo-electron tomography

Cryo-electron tomography was performed as previously described (32), with modifications as described below.

#### EM Grid Preparation and Cell Seeding

Gold quantifoil R2/2 200 mesh EM grids (EMS catalog: Q250-AR2) were glow-discharged at 25 mA for 30 seconds. Grids were then floated on drops of 0.01% poly-l-lysine (Sigma catalog: A-005-M) in a laminar flow hood for 30 min. Eight-chamber slides with removable wells (Sigma, PEZGS0816) were coated with 250 μL of 0.01% poly-l-lysine (Sigma, A-005-M) per well. U2OS cells transfected with sgSUV3 were trypsinized using 0.25% trypsin (Fisher Scientific, 25-200-056). Poly-l-lysine was removed from the wells and replaced with fresh DMEM, and EM grids were rinsed in DMEM prior to insertion into the bottom of the well of the eight-chamber plate. The cells were resuspended at 100,000 cells/mL, and 200 μL of the cell suspension (∼20,000 cells per grid) was added at the top of each EM grid in the wells. Cells were allowed to attach and recover overnight at standard culture conditions.

#### Plunge Freezing and Cryo-FIB Milling

Grids were retrieved, rinsed three times with phosphate-buffered saline (PBS), and back-blotted using a filter paper and a Teflon sheet using a Vitrobot Mark IV (ThermoFisher) equilibrated to 90% humidity and 37°C. Grids were plunge-frozen into liquid ethane and then clipped onto notched bases for cryo-FIB milling (ThermoFisher).Clipped grids were loaded into an Aquilos 2 (ThermoFisher) and candidate lamellae sites were screened via SEM. Grids were sputter-coated for 15 seconds at 30 mA and 10 Pa, followed by 1 minute of gas injection system (GIS) coating. Lamellae were generated using Auto-TEM: initial ablation at 0.3-0.5 nA to reach ablate the material to 3 μm thickness, thinning at 100 pA to 1 μm, 50 pA to thin cells to 500 nm, and final thinning at 30 pA to achieve 200 nm thickness.

#### Cryo-ET Data Acquisition

Lamellae-containing grids were immediately loaded into a 300kV Titon Krios microscope (Thermo Fisher Scientific). The autogrids containing lamellae were loaded such that the pretilt axis induced by FIB milling was perpendicular to the tilt axis of the microscope. Montage maps were generated for the entire autogrid to identify lamellae positions and a second medium mag montage generated at each lamellae site. Polygon montages were used to outline the borders of each lamella and used to guide data collection. Tilt series were collected using a dose-symmetric scheme starting from 10 to 15° lamellae pretilt with increments of 3° in groups of two tilts. The nominal defocus was varied between tilt series from-2 to-6 µm with a step size of 0.25 µm. The total dose per tilt series was approximately 120 e^-^/Å^2^.

#### Tomogram Reconstruction and Analysis

Scipion3 was used to facilitate software packages for downstream processing. Tilt series were corrected for beam-induced motion and binned in Fourier space using MotionCor3 (RRID:SCR_01649). CTF estimation was performed with CTFfind5, and tilt series alignment and tomogram reconstruction were performed in AreTomo2. Tomograms were denoised using IsoNet2 (55) (https://github.com/IsoNet-cryoET/IsoNet2) and segmented using Membrain with default parameters. Cristae surface area and volume were measured in ChimeraX, and statistical significance was assessed with unpaired t-tests in GraphPad Prism 10. Dragonfly (RRID:SCR_025150) was used to manually segment condensates and calcium clusters using the multi-slice brush and local otsu method for visualization only.

#### ATP Synthase Subtomogram Averaging and Template Matching

ATP synthase complexes were manually picked in Napari and imported into Relion 5 for subtomogram averaging. Initial models were generated and refined, followed by template matching in PyTOM using a cross-correlation score of 0.44 and membrane-based masks expanded by 100 Å. EM density maps were visualized in ChimeraX, and differences in ATP synthase localization were assessed using Fisher’s Exact test.

### Statistical analyses

Following analysis, the data values were exported to.xlsx files, and all data organization and calculations were performed in Microsoft Excel. Statistical analyses and plotting of the data were done using GraphPad Prism (version 11.0.0). Please refer to the figure legends or the sub-sections of Material and Methods for details on number of biological replicates (N) and statistical details. For experiments in which two groups were compared, two-tailed Mann-Whitney test was used except otherwise noted. Error bars display mean values ± standard deviation. P-values under 0.05 were considered significant.

### Illustrations

Illustrations were created using Adobe Illustrator.

## ACKNOWLEDGMENTS

We thank the members of the Lewis laboratory and the ASAP team mito911 for helpful discussions and advice. We thank Dr. Adam Begeman and Hera Saqub for assistance with sample preparation and microscopy.

This research was funded by Aligning Science Across Parkinson’s [ASAP-000350] through the Michael J. Fox Foundation for Parkinson’s Research (MJFF) (J.H.H, and S.C.L.). A portion of this work was supported by the National Institutes of Health grant R35GM147218 (S.C.L.), and the University of California Cancer Research Coordinating Committee grant C27DF11281 (E.V.K.). Some of this work was performed at the Stanford-SLAC CryoET Specimen Preparation Center (SCSC), which is supported by the National Institutes of Health Common Fund’s Transformative High Resolution Cryoelectron Microscopy program (U24GM139166).

## DATA AVAILABILITY STATEMENT

All information needed to evaluate the conclusions in the paper are present in the paper and/or the supplementary materials. Data, key material and code are listed in a Key Resource Table at supplementary table S1, and will be available via the BioStudies data repository prior to or upon acceptance for publication. Please see the Lewis laboratory page at protocols.io, as an additional resource.

## COMPETING INTERESTS

JHH is a shareholder of Casma Therapeutics.

