## Supplemental Figures S1-S9 for "Nanoscale remodeling of mitochondrial architecture during mtRNA surveillance failure"

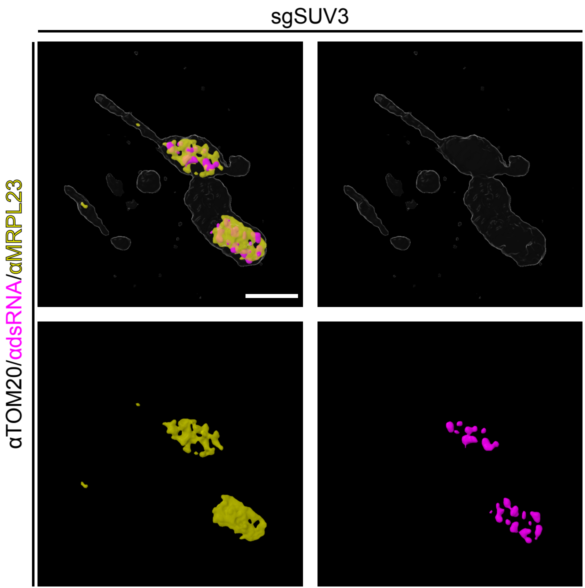
sgSUV3
αTOM20/αdsRNA/αMRPL23

**Figure S1: Mitochondrial organization of MRPL23 and dsRNA within stress bodies.**
3D rendering of the mitochondria shown in image (i.) from figure 1C. Scale bar: 2μm.

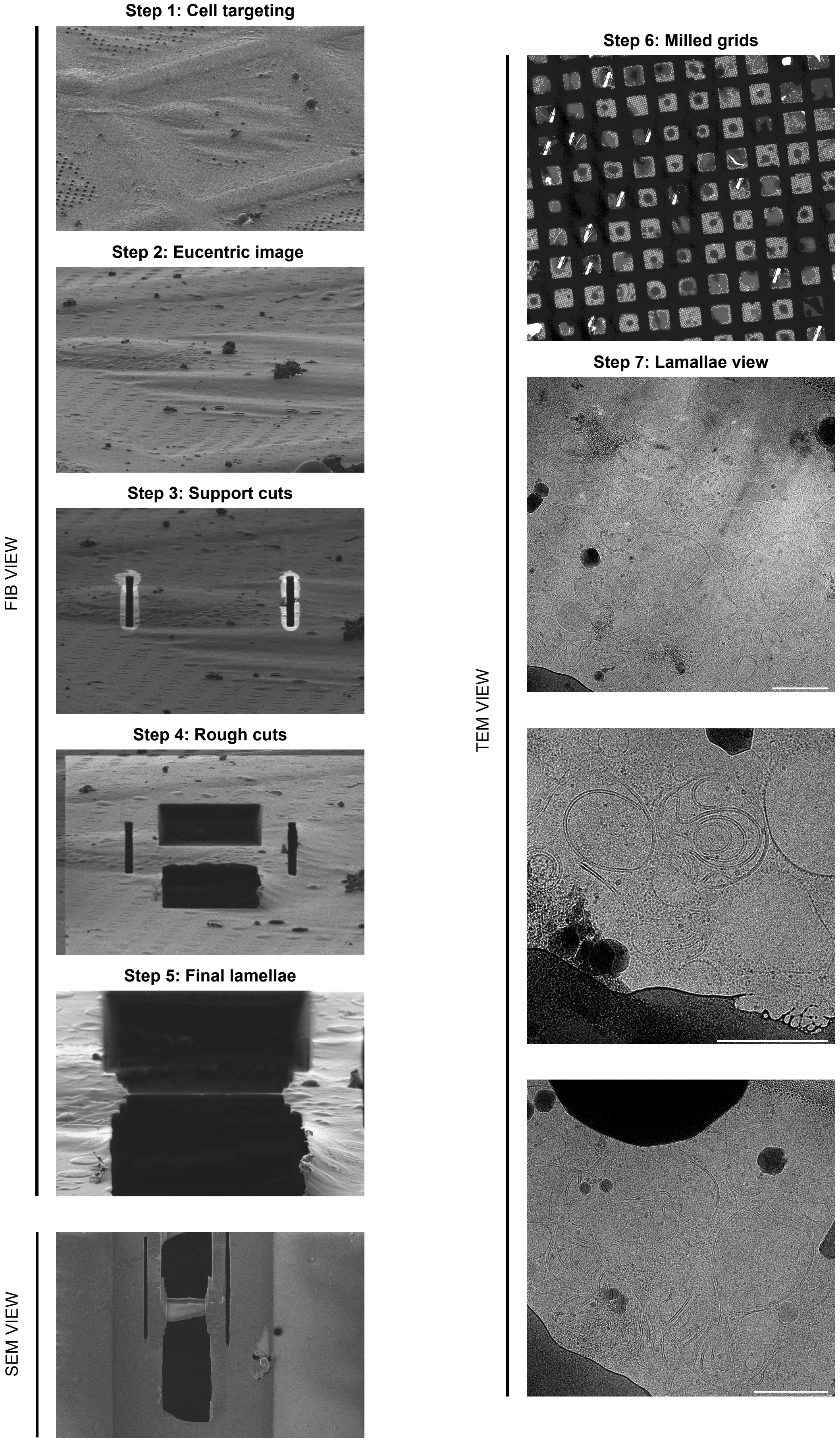
Step 1: Cell targeting
Step 2: Eucentric image
Step 3: Support cuts
Step 4: Rough cuts
Step 5: Final lamellae
FIB VIEW
SEM VIEW
Step 6: Milled grids
Step 7: Lamallae view
TEM VIEW

**Figure S2: Process of acquiring cryo-ET tomograms.**
Distinct steps of acquisition of the tomograms in this study. Scale bar: 1µm.

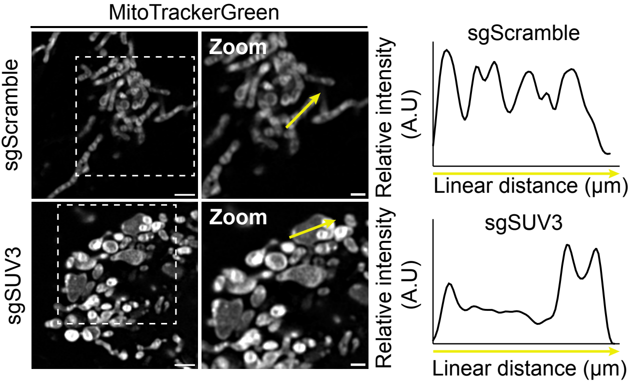
MitoTrackerGreen
sgScramble
Zoom
Relative intensity (A.U)
sgScramble
Linear distance (µm)
sgSUV3
Zoom
Relative intensity (A.U)
sgSUV3
Linear distance (µm)

**Figure S3: SUV3 depletion induces inner mitochondrial membrane remodeling in live cells.** Left: live imaging of MitoTrackerGreen in U2OS-Cas9 transfected with sgScramble (top) or sgSUV3 (bottom). Scale bars: 2 µm; 1 µm in zoom. A single slice of a z-stack is represented for each condition. Right: Relative MitoTracker Green intensity plots for the yellow arrow drawn on the left.

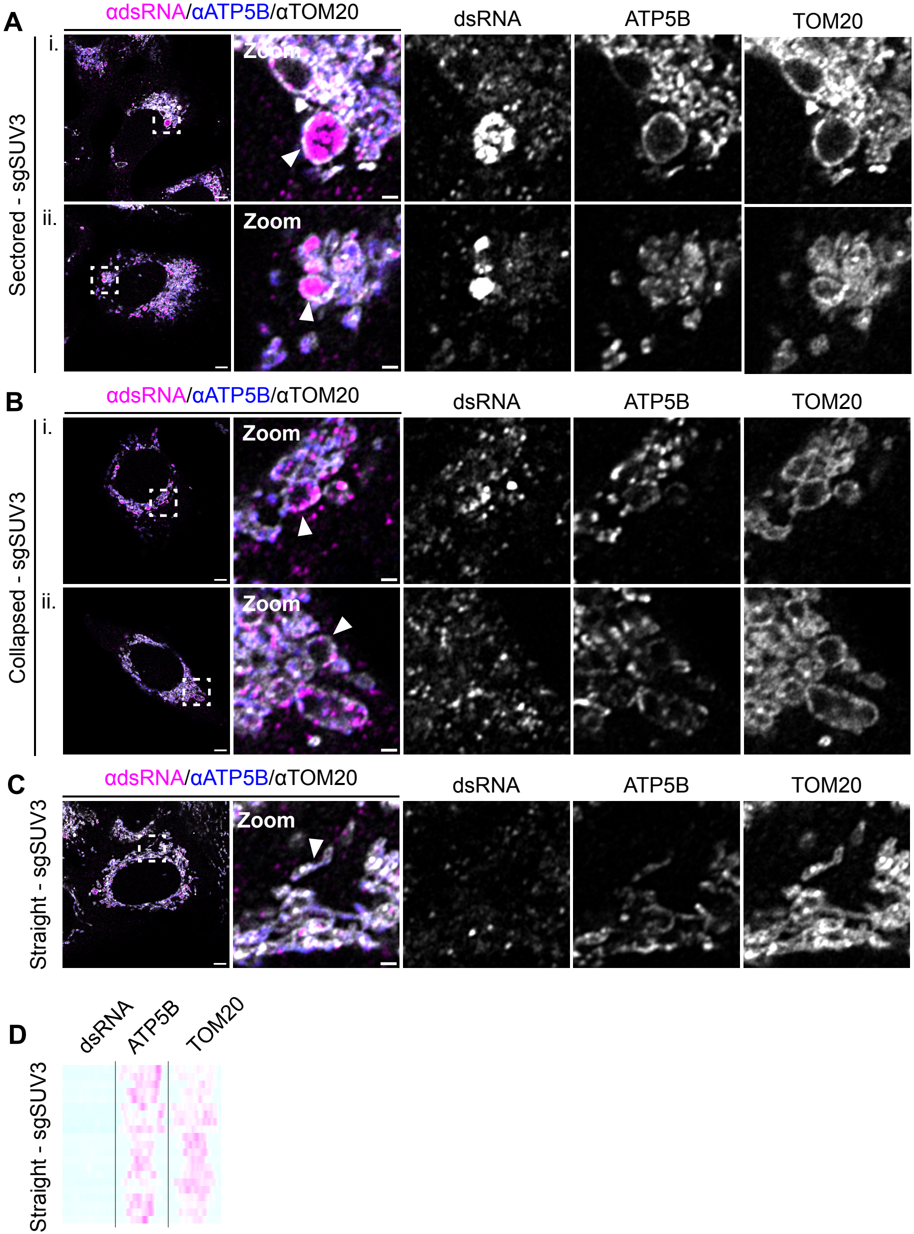

A
αdsRNA/αATP5B/αTOM20
dsRNA
ATP5B
TOM20
Sectored - sgSUV3
i.
ii.
Zoom
B
Collapsed - sgSUV3
C
Straight - sgSUV3
D

**Figure S4: Mitochondrial phenotype variety in sgSUV3 cells.**
Immunofluorescence images of U2OS-Cas9 transfected with sgSUV3 and immunolabeled with anti-TOM20, anti-dsRNA (J2), and anti-ATP5B. Imaged 7 days after transfection. **(A)** Additional examples of mitochondria with sectored dsRNA in sgSUV3-cells. **(B)** Additional examples of mitochondria with collapsed dsRNA in sgSUV3 cells. **(C)** Example of straight mitochondria observed in sgSUV3 cells. A single slice of a z-stack is shown for all images represented. Scale bars: 5 μm; 1 μm in zoom. Arrows point towards mitochondria that have been used for the heatmap analysis (F4B, FS4D), and images shown come from any of the three biological replicates used in the heatmap analysis. (**D**) Relative intensity heatmaps of TOM20, dsRNA, and ATP5B for straight mitochondria observed in sgSUV3. 21 mitochondria were analyzed across three biological replicates with 7 mitochondria analyzed per replicate.

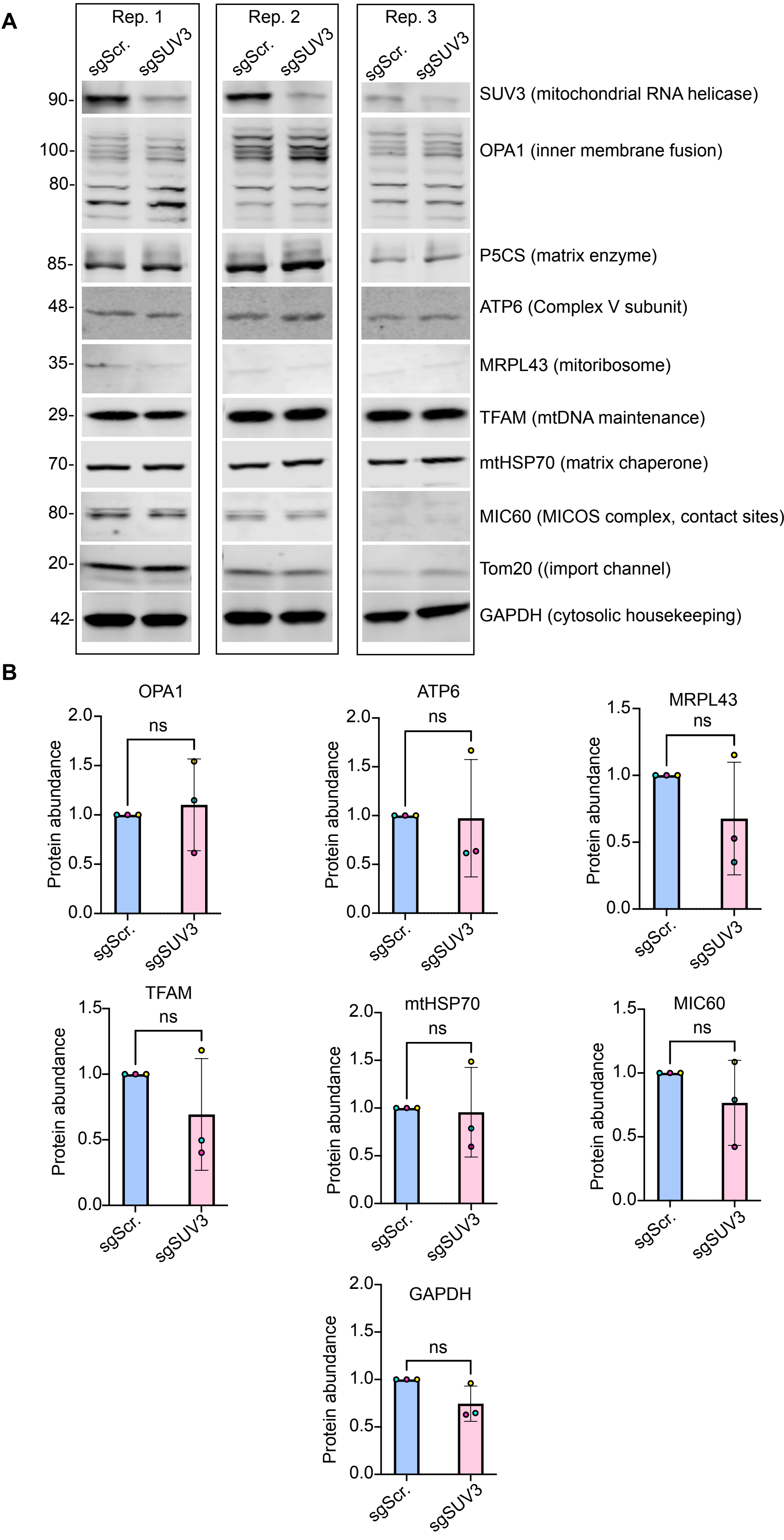
A
Rep. 1
Rep. 2
Rep. 3
sgScr.
sgSUV3
90-
100-
80-
85-
48-
35-
29-
70-
80-
20-
42-
SUV3 (mitochondrial RNA helicase)
OPA1 (inner membrane fusion)
P5CS (matrix enzyme)
ATP6 (Complex V subunit)
MRPL43 (mitoribosome)
TFAM (mtDNA maintenance)
mtHSP70 (matrix chaperone)
MIC60 (MICOS complex, contact sites)
Tom20 ((import channel)
GAPDH (cytosolic housekeeping)
B
OPA1
ATP6
MRPL43
TFAM
mtHSP70
MIC60
GAPDH
ns
Protein abundance
0.0
0.5
1.0
1.5
2.0
sgScr.
sgSUV3

**Figure S5: Protein levels of mitochondrial proteins of interest following SUV3 depletion.** **(A)** Levels of proteins of interest in sgScramble versus sgSUV3 U2OS-Cas9 cells analyzed by western blot. GAPDH as the loading control. **(B)** Quantification of the blots represented in (A) normalized to TOM20 and represented as a fold change relative to sgScramble. Error bars display mean values ± standard deviation. Statistical significance determined with an unpaired t-test. N = 3 independent biological replicates.

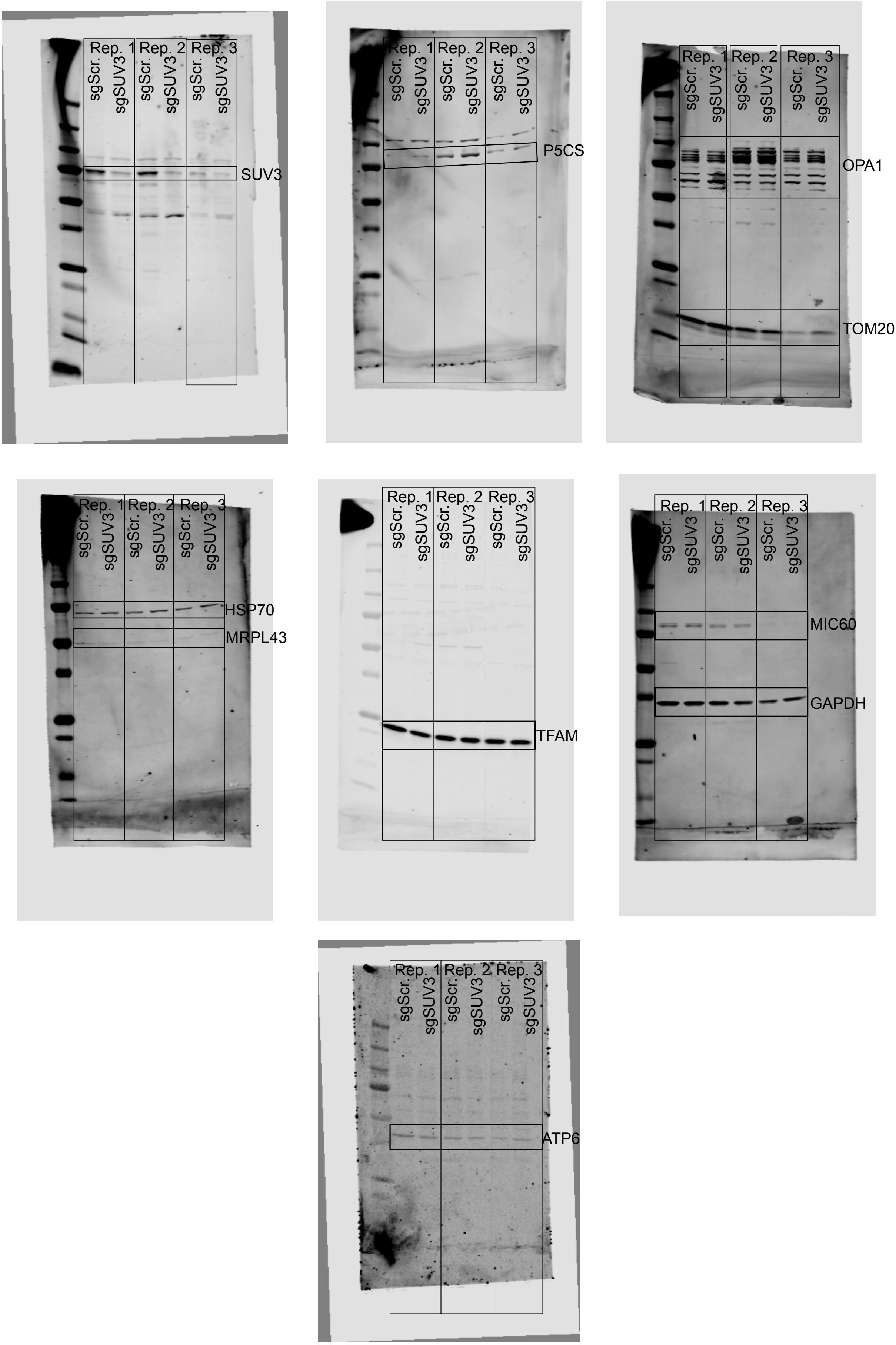

Rep. 1 Rep. 2 Rep. 3
sgScr. sgSUV3 sgScr. sgSUV3 sgScr. sgSUV3
SUV3
P5CS
OPA1
TOM20
HSP70
MRPL43
TFAM
MIC60
GAPDH
ATP6

**Figure S6: Uncropped western blots.**
Uncropped full field of view of the western blots represented in Fig. S5.

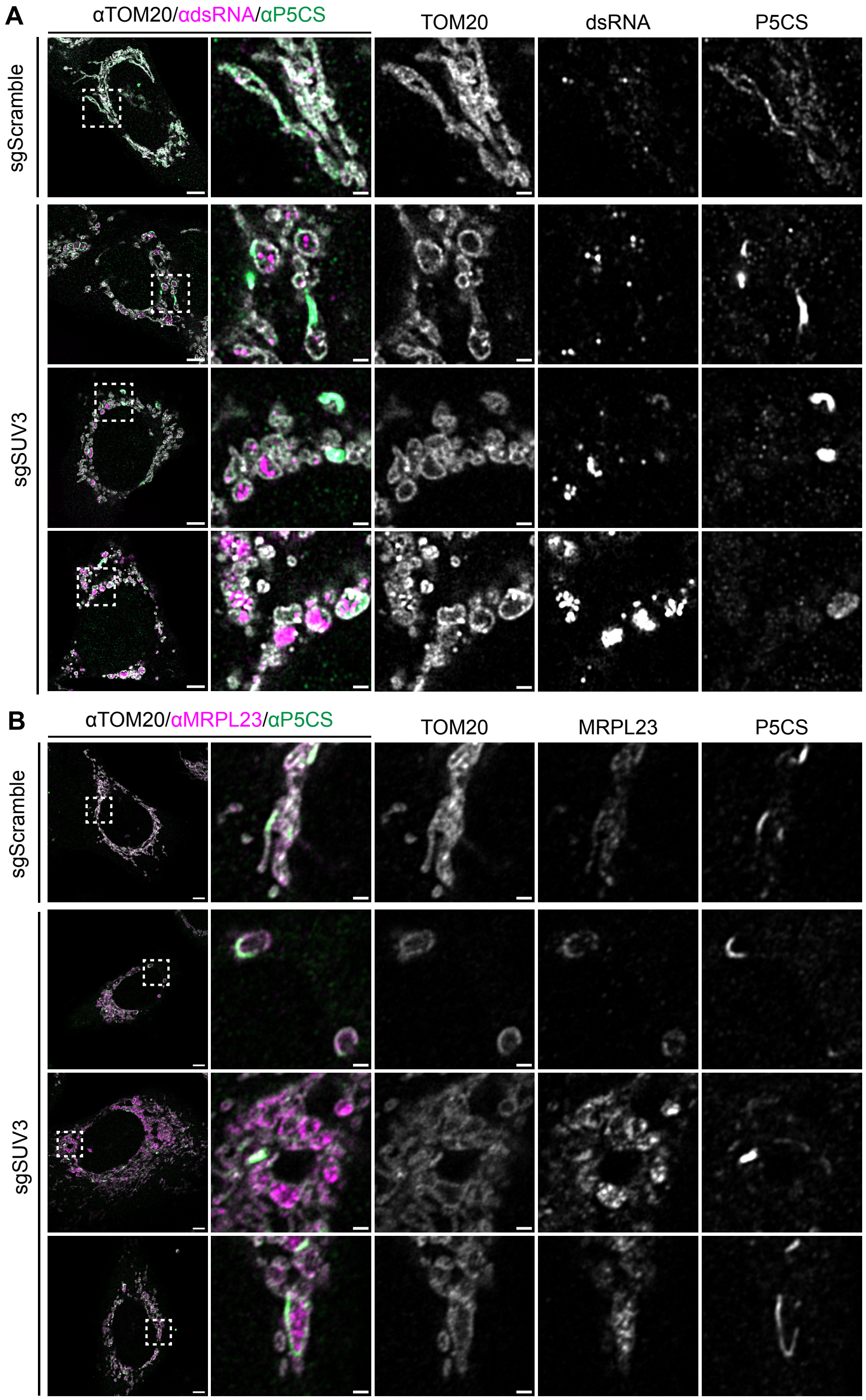
A
αTOM20/αdsRNA/αP5CS
TOM20
dsRNA
P5CS
sgScramble
sgSUV3
B
αTOM20/αMRPL23/αP5CS
TOM20
MRPL23
P5CS
sgScramble
sgSUV3

**Figure S7: Localization of P5CS and MSB components following SUV3 depletion.**
**(A)** Enlarged images of the individual channels shown in figure 5G. **(B)** Enlarged images of the individual channels shown in figure 5H. Scale bars: 5μm (full image); 1μm (zoom).

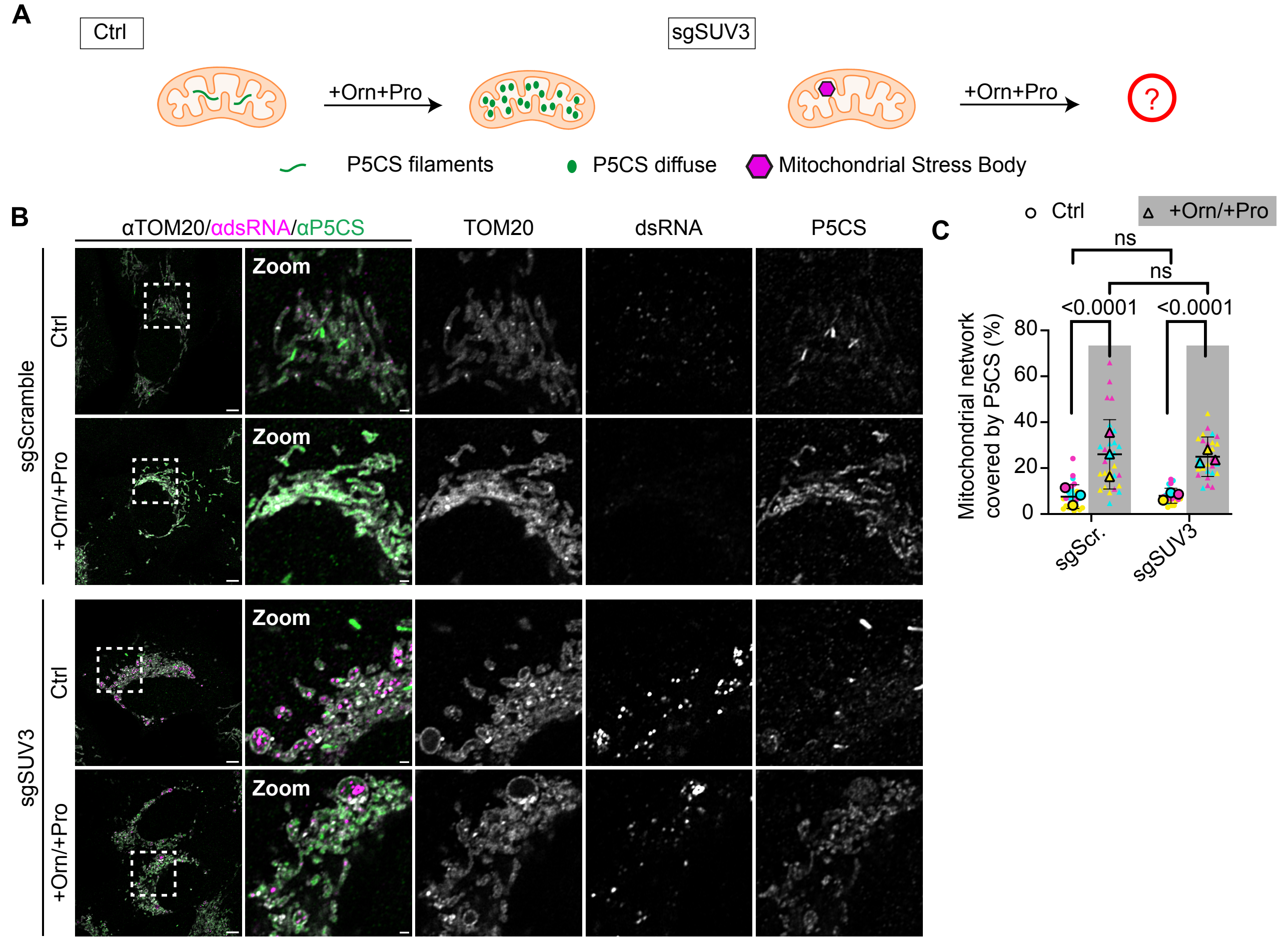
A
Ctrl
+Orn+Pro
sgSUV3
+Orn+Pro
?
P5CS filaments
P5CS diffuse
Mitochondrial Stress Body
B
αTOM20/αdsRNA/αP5CS
TOM20
dsRNA
P5CS
sgScramble
Ctrl
+Orn/+Pro
sgSUV3
Ctrl
+Orn/+Pro
Zoom
C
Ctrl
+Orn/+Pro
ns
ns
<0.0001
<0.0001
Mitochondrial network covered by P5CS (%)
0
20
40
60
80
sgScr.
sgSUV3

**Figure S8: Ornithine/Proline supplementation does not affect dsRNA distribution.**
**(A)** Combined ornithine/proline supplementation results in redistribution of P5CS along the mitochondrial network in control cells, but has unknown effects on dsRNA distribution in sgSUV3 cells. **(B)** Immunofluorescence images of U2OS-Cas9 transfected with sgScramble (top) with (+Orn/+Pro) or without (control) ornithine/proline supplementation and sgSUV3 (bottom) with (+Orn/+Pro) or without (control) ornithine/proline supplementation. Cells were stained with anti-TOM20, anti-dsRNA (J2), and anti-P5CS and imaged 7 days after transfection. Scale bars: 5 μm; 1 μm in zoom. A single slice of a z-stack is shown for each condition. **(C)** Quantification of total mitochondrial P5CS area covering the mitochondrial network performed on maximum intensity projection images from the experiment in (B). N = 28 FOV (sgScramble; control), 30 FOV (sgScramble; +Orn/+Pro), 27 FOV (sgSUV3; control), 27 FOV (sgSUV3; +Orn/+Pro). Error bars display mean values ± standard deviation, and bottom bar is clipped if below 0. Statistical analysis performed with Mann-Whitney test on individual data points.

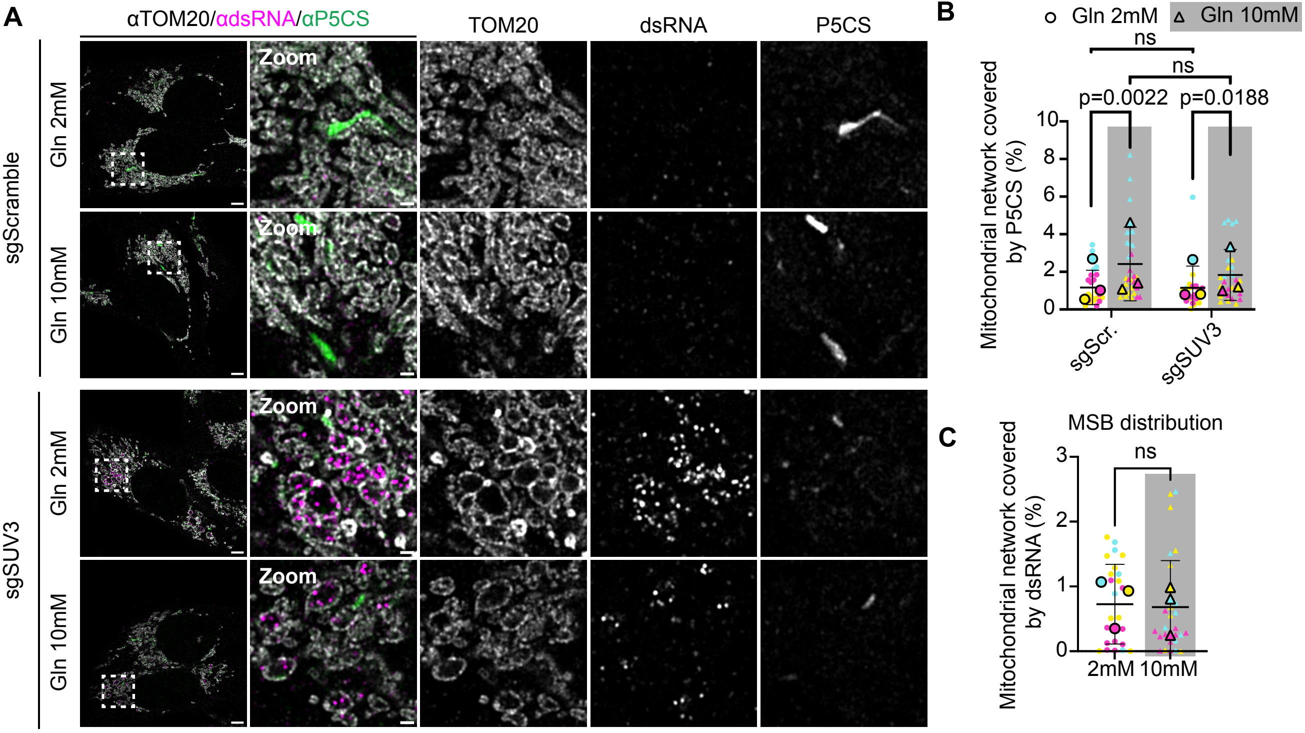
A
αTOM20/αdsRNA/αP5CS
TOM20
dsRNA
P5CS
sgScramble
sgSUV3
Gln 2mM
Gln 10mM
Zoom
B
Gln 2mM
Gln 10mM
ns
ns
p=0.0022
p=0.0188
Mitochondrial network covered by P5CS (%)
sgScr.
sgSUV3
C
MSB distribution
ns
Mitochondrial network covered by dsRNA (%)
2mM
10mM

**Figure S9: Glutamine supplementation does not impact dsRNA distribution.**
**(A)** Immunofluorescence images of U2OS-Cas9 transfected with sgScramble (top) grown with 2mM or 10mM glutamine (Gln) and sgSUV3 (bottom) grown with 2mM or 10mM Gln. Cells were stained with anti-TOM20, anti-dsRNA(J2), and anti-P5CS and imaged 7 days after transfection. Scale bars: 5 µm; 1 µm in zoom. A single slice of a z-stack is shown for each condition. **(B)** Quantification of total mitochondrial P5CS area covering the mitochondrial network. Quantification performed on maximum intensity projection images from experiment in (A). **(C)** Quantification of total mitochondrial dsRNA area covering the mitochondrial network in sgSUV3 cells. Quantification performed on maximum intensity projection images from experiment in (A). For (B), N = 23 FOV (sgScramble, 2mM Gln); N = 30 FOV (sgScramble, 10mM Gln); N =24 FOV (sgSUV3, 2mM Gln); N = 29 FOV (sgSUV3, 10mM Gln). For (C), N = 25 FOV (sgSUV3, 2mM Gln); N = 30 FOV (sgSUV3, 10mM Gln) across three independent biological replicates. Error bars display mean values ± standard deviation, and the bottom bar is clipped if below 0. Statistical analysis performed with Mann-Whitney test on individual data points.
