## Supplementary material for "Nanoscale remodeling of mitochondrial architecture during mtRNA surveillance failure": Table S2

|  | **Guide RNAs** | | |
| --- | --- | --- | --- |
|  | **guide_#1 (5'-3')** | **guide_#2 (5'-3')** | **guide_#3 (5'-3')** |
| **SUPV3L1** | CCUAUUGUGGGCUCGGCUCC | GGGCCCUUUCCCGGGGUUCU | UUUGGAGCCACCGGAGGCAG |
| **Non-targeting control 1** | GCACUACCAGAGCUAACUCA | N/A | N/A |
| **Non-targeting control 2** | GUACGUCGGUAUAACUCCUC | N/A | N/A |

**Table S2**. **Sequences of guide RNAs for the generation of CRISPR knockouts.** Non-targeting control 1 (EditCo) was used as the control in most experiments. In Figure S8, non-targeting control 2 (EditCo) was used instead.
