## Supplementary material for "Nanoscale remodeling of mitochondrial architecture during mtRNA surveillance failure": Table S3

| **Name** | **5'-3' Seq** |
| --- | --- |
| **SUPV3L1_PCR_Forward** | ATACTCGTACCCTGAGCATTC |
| **SUPV3L1_PCR_Reverse** | GGCGTCCACCTAACAACTC |
| **SUPV3L1_Sequencing_Forward** | AGCTTACCTGCTCGGCCT |
| **SUPV3L1_Sequencing_Reverse** | CTTCCTCCCAACCAGCCTC |

**Table S3**. **Sequences of primers used for CRISPR knockout validation**
