## Supplementary material for "Nanoscale remodeling of mitochondrial architecture during mtRNA surveillance failure": Table S4

|  | Rep1 | Rep2 | Rep3 | Rep4 | Rep5 | Rep6 | Rep7 | Average |
| --- | --- | --- | --- | --- | --- | --- | --- | --- |
| SUPV3L1 KO Efficiency (%) | 33.5 | 28.5 | 37.5 | 70.5 | 73 | 71.5 | 76.5 | 55.9 |

**Table S4**. **ICE % KO efficiency across seven experimental replicates.** Non-targeting control 1 was used as a control sample for sequence comparison. %KO Efficiency is reported as the average of scores obtained from the forward sequence and reverse sequence.
